# Synthetic Multiantigen MVA Vaccine COH04S1 Protects Against SARS-CoV-2 in Syrian Hamsters and Non-Human Primates

**DOI:** 10.1101/2021.09.15.460487

**Authors:** Flavia Chiuppesi, Vu H. Nguyen, Yoonsuh Park, Heidi Contreras, Veronica Karpinski, Katelyn Faircloth, Jenny Nguyen, Mindy Kha, Daisy Johnson, Joy Martinez, Angelina Iniguez, Qiao Zhou, Teodora Kaltcheva, Paul Frankel, Swagata Kar, Ankur Sharma, Hanne Andersen, Mark G. Lewis, Yuriy Shostak, Felix Wussow, Don J. Diamond

**Author notes:** These authors jointly supervised this work (Felix Wussow, Don J. Diamond).

## Abstract

Second-generation COVID-19 vaccines could contribute to establish protective immunity against SARS-CoV-2 and its emerging variants. We developed COH04S1, a synthetic multiantigen Modified Vaccinia Ankara-based SARS-CoV-2 vaccine that co-expresses spike and nucleocapsid antigens. Here, we report COH04S1 vaccine efficacy in animal models. We demonstrate that intramuscular or intranasal vaccination of Syrian hamsters with COH04S1 induces robust Th1-biased antigen-specific humoral immunity and cross-neutralizing antibodies (NAb) and protects against weight loss, lower respiratory tract infection, and lung injury following intranasal SARS-CoV-2 challenge. Moreover, we demonstrate that single-dose or two-dose vaccination of non-human primates with COH04S1 induces robust antigen-specific binding antibodies, NAb, and Th1-biased T cells, protects against both upper and lower respiratory tract infection following intranasal/intratracheal SARS-CoV-2 challenge, and triggers potent post-challenge anamnestic antiviral responses. These results demonstrate COH04S1-mediated vaccine protection in animal models through different vaccination routes and dose regimens, complementing ongoing investigation of this multiantigen SARS-CoV-2 vaccine in clinical trials.

## Introduction

Since the emergence of severe acute respiratory syndrome coronavirus 2 (SARS-CoV-2) in China at the end of 2019, SARS-CoV-2 has spread rapidly worldwide, causing a global pandemic with millions of fatalities^1,2^. Several SARS-CoV-2 vaccines were developed in response to the COVID-19 pandemic with unprecedented pace and showed 62-95% efficacy in Phase 3 clinical trials, leading to their emergency use authorization (EUA) in many countries by the end of 2020/beginning of 2021^3–8^. This includes vaccines based on mRNA, adenovirus vectors, and nanoparticles that utilize different antigenic forms of the spike (S) protein to induce protective immunity against SARS-CoV-2 primarily through the function of neutralizing antibodies (NAb)^9–12^. While the Phase 3 efficacy results provide hope for a rapid end of the COVID-19 pandemic, waning antibody responses and evasion of NAb by emerging variants of concern (VOC) pose imminent challenges for durable vaccine protection and herd immunity^13–19^. Alternative vaccines based on different platforms or modified epitope or antigen design could therefore contribute to establish long-term and cross-reactive immunity against SARS-CoV-2 and its emerging VOC.

While NAb blocking S-mediated entry are considered the principal SARS-CoV-2 immune correlate of protection, humoral and cellular immune responses to multiple antigens have been implicated in the protection against SARS-CoV-2^10,11,20^. Besides the S protein, the nucleocapsid (N) protein is well-recognized as a dominant target of antibody and T cell responses in SARS-CoV-2-infected individuals and therefore suggested as an additional immunogen to augment vaccine-mediated protective immunity^21–24^. Its high conservation and universal cytoplasmic expression makes the N protein an attractive complementary target antigen to elicit durable and broadly reactive T cells^23^. Several recent studies highlight the benefits of N as a vaccine antigen in animal models^25–28^.

We previously constructed multiantigenic SARS-CoV-2 vaccine candidates using a fully synthetic platform based on the well-characterized and clinically proven Modified Vaccinia Ankara (MVA) vector^29–32^, which is marketed in the USA as Jynneos™ (Bavarian-Nordic)^33^. MVA is a highly attenuated poxvirus vector that is widely used to develop vaccines for infectious diseases and cancer due to its excellent safety profile in animals and humans, versatile delivery and expression system, and ability to stimulate potent humoral and cellular immune responses to heterologous antigens^30–32^. We have used MVA to develop vaccine candidates for preclinical testing in animal models of congenital cytomegalovirus disease while demonstrating vaccine efficacy in several clinical trials in solid tumor and stem cell transplant patients^34–39^. Using the synthetic MVA (sMVA) platform, we constructed sMVA vectors co-expressing full-length S and N antigen sequences that showed potent immunogenicity in mice to stimulate SARS-CoV-2 antigen-specific humoral and cellular immune responses, including high-titer NAb^29^. One of these sMVA constructs forms the basis of clinical vaccine candidate COH04S1, which has shown to be safe and immunogenic in a randomized, double-blinded, placebo-controlled, single center Phase 1 trial in healthy adults (NCT04639466), and is currently evaluated in a randomized, double-blinded, single center Phase 2 trial in hematology patients who have received cellular therapy (NCT04977024).

In this report, we demonstrate that COH04S1 stimulates protective immunity against SARS-CoV-2 in Syrian hamsters by intramuscular (IM) and intranasal (IN) vaccination and in non-human primates (NHP) through two-dose (2D) and single-dose (1D) vaccination regimens. These results complement the clinical evaluation of this multiantigen sMVA-based SARS-CoV-2 vaccine.

## Results

### COH04S1 induces robust Th1-biased S and N antigen-specific antibodies and cross-NAb against SARS-CoV-2 in hamsters through IM and IN vaccination

Syrian hamsters are widely used to evaluate vaccine protection against SARS-CoV-2 in a small animal model that mimics moderate-to-severe COVID-19 disease^40–45^. Using this animal model, we assessed efficacy of COH04S1 (Fig. 1a) to protect against SARS-CoV-2 challenge by either IM or IN vaccination to assess vaccine protection via parenteral or mucosal immune stimulation. Hamsters were vaccinated twice in a four week interval with 1×10^8^ plaque forming units (PFU) of COH04S1 by IM or IN route, referred to as COH04S1-IM or COH04S1-IN, respectively (Fig. 1b). Unvaccinated animals and hamsters vaccinated IM or IN with empty sMVA vector were used as controls. COH04S1-IM and COH04S1-IN stimulated robust and comparable binding antibodies to both the S and N antigens, including antibodies to the S receptor binding domain (RBD), the primary target of NAb^46–48^. Binding antibodies to S, RBD, and N were detected at high levels in both the COH04S1-IM and COH04S1-IN vaccine groups after the first vaccination and boosted after the second vaccination (Figs. 1c-d and S1), whereby a particularly strong booster effect was observed for RBD-specific antibodies. Antigen-specific binding antibodies stimulated by COH04S1-IM and COH04S1-IN in hamsters were mainly composed of IgG2/3 isotypes and only to a minor extent of IgG1 isotype (Figs. 1e and S1), indicating Th1-biased immune responses.

**Figure 1.**
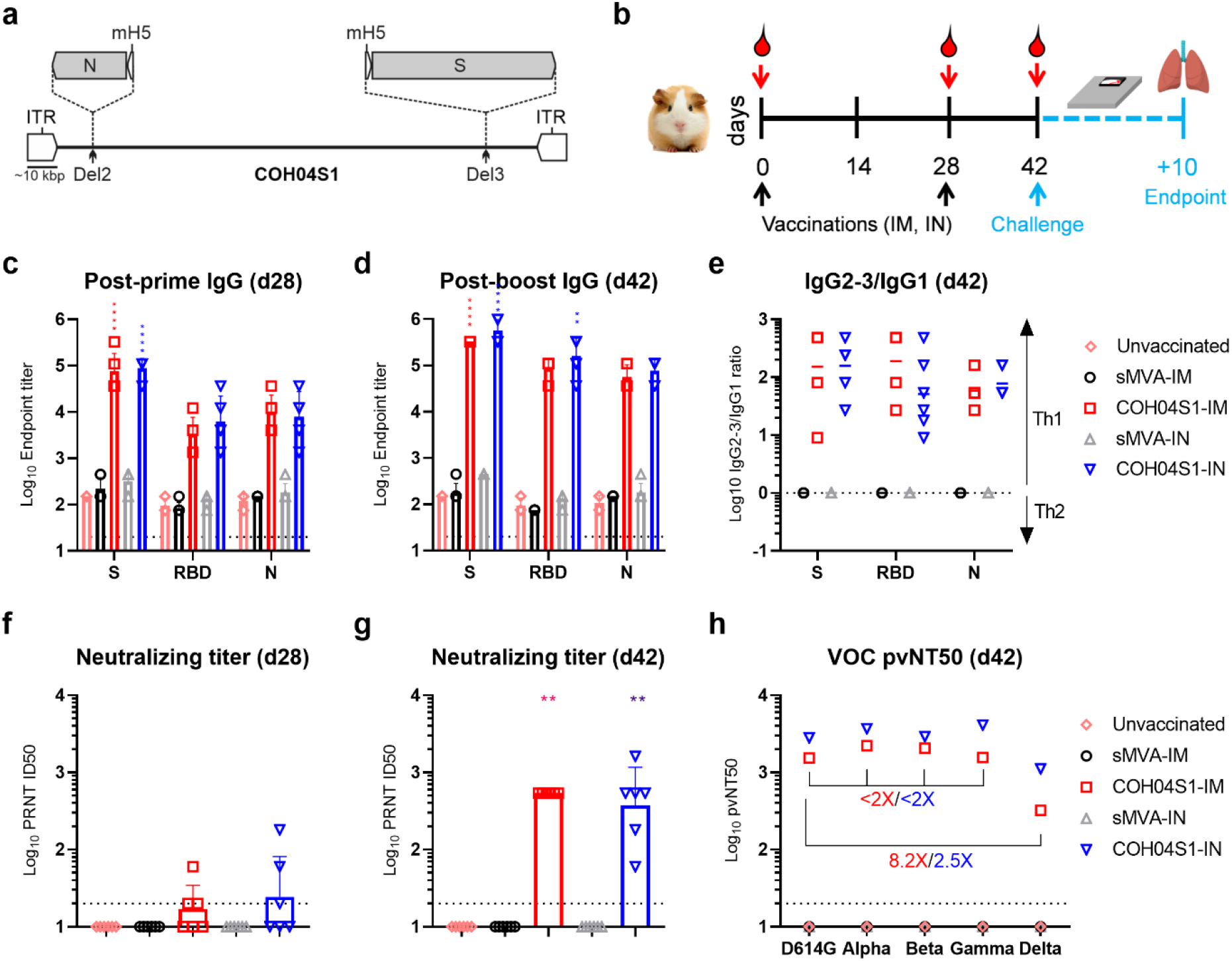
COH04S1 immunogenicity in Syrian hamsters. **a. COH04S1 construct.** The sMVA-based COH04S1 vaccine vector co-expresses SARS-CoV-2 S and N antigen sequences that are inserted together with mH5 promoter elements into the MVA Deletion 2 (Del2) and Deletion 3 (Del3) sites as indicated. ITR = inverted terminal repeat. **b. Study design.** Hamsters were vaccinated twice with COH04S1 by IM (COH04S1-IM) or IN (COH04S1-IN) route as indicated (black arrows). Unvaccinated animals and hamsters vaccinated with empty sMVA vector by IM (sMVA-IM) or IN (sMVA-IN) route were used as controls. Blood samples were collected at 0, 28, and 42 day (red arrows). Hamsters were challenged IN at day 42 and body weight changes were recorded daily for 10 days. At endpoint, nasal wash, turbinates and lung tissue were collected for downstream analyses. **c-d. IgG endpoint titers.** S, RBD, and N antigen-specific binding antibody titers were measured in serum samples of vaccine and control groups at day 28 (d28) post prime and at day 42 (d42) post booster immunization via ELISA. Data are presented as geometric mean values + geometric SD. Dotted lines indicate lower limit of detection. Two-way ANOVA with Tukey’s multiple comparison test was used. **e. IgG2-3/IgG1 ratios**. S, RBD, and N antigen-specific IgG2-3 and IgG1 endpoint titers were measured at day 42 (d42) in serum samples of vaccine and control groups and used to assess IgG2-3/IgG1 ratios. An IgG2-3/IgG1 ratio >1 is indicative of a Th1-biased response. Geometric means are indicated with a line. **f-g. NAb titers.** NAb titers were measured in serum samples of vaccine and control groups post-first (d28) and post-second (d42) immunization via PRNT assay against SARS-CoV-2 infectious virus. Data are presented as geometric mean values + SD. Dotted lines indicate lower limit of detection. Values below the limit of detection (PRNT=20) are indicated as 10. One-way ANOVA with Holm-Sidak’s multiple comparison test was used. **h. VOC-specific NAb titers.** NAb were measured in pooled serum samples of vaccine and control groups collected at the time of challenge (d42) using pseudovirus (pv) variants with S D614G mutation or S modifications based on several VOC, including Alpha (B.1.1.7), Beta (B.1.351), Gamma (P.1), and Delta (B.1.617.2). Titers are expressed as NT50. Fold NT50 titer reduction in comparison to D614G PV are shown. *0.05 < p <0.01, **0.01 < p < 0.001, ***0.001 < p < 0.0001, ****p < 0.0001.

Plaque reduction neutralization titer (PRNT) assay measuring neutralizing activity against SARS-CoV-2 infectious virus (USA-WA1/2020) demonstrated that potent and comparable NAb titers were stimulated by COH04S1-IM and COH04S1-IN after the booster vaccinations (Figs. 1f-g). Furthermore, pooled post-boost immune sera from COH04S1-IM and COH04S1-IN-vaccinated animals demonstrated potent cross-reactive neutralizing activity against SARS-CoV-2 pseudoviruses (PV) with D614G S mutation or multiple S modifications based on several SARS-CoV-2 VOC (Figs. 1h and S1 and Table S1), including Alpha (B.1.1.7), Beta (B.1.351), Gamma (P.1), and the increasingly dominant Delta variant (B.1.617.2). Notably, while NAb titers measured with Alpha, Beta, and Gamma PV variants were generally similar to those determined with D614G PV, NAb titers measured with the Delta-matched PV variant were 2-8-fold reduced compared to those determined with D614G PV. In addition, NAb titers measured in COH04S1-IM immune sera using the different PV variants appeared overall lower than those measured in COH04S1-IN immune sera. These results demonstrate that IM and IN vaccination of Syrian hamsters with COH04S1 elicits robust Th1-biased S and N antigen-specific binding antibodies as well as potent NAb responses with cross-reactivity to prevent PV infection by several SARS-CoV-2 VOC.

### IM and IN vaccination with COH04S1 protect hamsters from progressive weight loss, lower respiratory tract infection, and lung pathology following SARS-CoV-2 challenge

Two weeks after the booster vaccination, hamsters were challenged IN with 3×10^4^ PFU of SARS-CoV-2 reference strain USA-WA1/2020 and body weight changes were measured over a period of 10 days. While control animals showed rapid body weight loss post challenge, hamsters vaccinated IM or IN with COH04S1 were protected from severe weight loss following challenge (Figs. 2a-b). Control animals showed rapid body weight loss for 6-7 days post challenge, reaching maximum weight loss between 10-20%. In contrast, COH04S1-IM and COH04S1-IN-vaccinated animals showed no or only very minor body weight decline post challenge, with maximum body weight loss below 4% for all animals at any time point during the entire 10 day observation period (Figs. 2a-b). Notably, while minor weight loss was observed for COH04S1-IM-vaccinated animals at 1-2 days post challenge, COH04S1-IN-vaccinated animals did not show body weight decline at these early time points post challenge (Fig. 2a), suggesting improved protection from weight loss by COH04S1 through IN compared to IM vaccination at an early phase post challenge.

**Figure 2.**
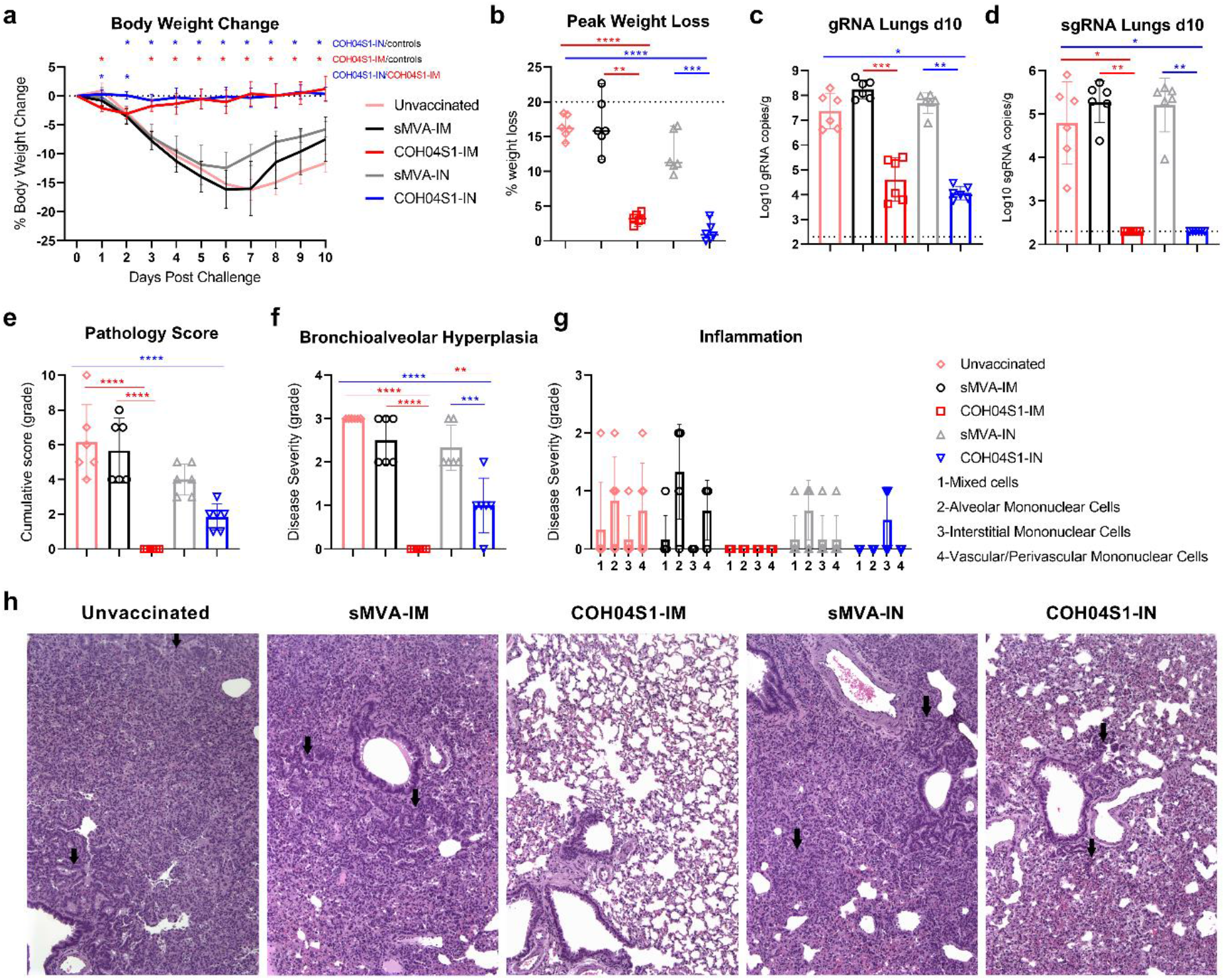
COH04S1-mediated vaccine protection in hamsters following sub-lethal SARS-CoV-2 challenge. **a. Body weight change.** Body weight of COH04S1-IM and COH04S1-IN-vaccinated animals as well as unvaccinated and sMVA-IM and sMVA-IN vector control animals was measured daily for 10 days post-challenge. Weight loss is reported as mean ±SD. Two-way ANOVA followed by Tukey’s multiple comparison test was used to compare group mean values at each timepoint. **b. Maximal weight loss.** Percentage of maximal weight loss is shown in single animals of vaccine and control groups. Lines and bars represent median values and 95% CI, respectively. Dotted line represents the maximum weight loss allowed before euthanasia. Peaks of weight loss in each group were compared using one-way ANOVA followed by Tukey’s multiple comparison test. **c-d. Lung viral loads.** SARS-CoV-2 genomic RNA (gRNA) and sub-genomic RNA (sgRNA) copies were quantified in lung tissue of vaccine and control groups at day 10 post-challenge by qPCR. Bars show RNA copies geometric mean ± geometric SD. Dotted lines represent lower limit of detection. Kruskal-Wallis test followed by Dunn’s multiple comparison test was used. **e-h. Histopathological findings.** Hematoxylin/eosin-stained lung sections of COH04S1-vaccinated hamsters and control animals at day 10 post challenge were evaluated by a board-certified pathologist and microscopic findings were graded based on severity on a scale from 1 to 5 (Table S2). Panel **e** shows the cumulative pathology score of all histopathologic findings in each group. Panel **f** shows grading of bronchioalveolar hyperplasia disease severity in each group. One-way ANOVA followed by Holm-Sidak’s multiple comparison test was used. Panel **g** shows the severity of lung inflammatory microscopic findings based on 1-to-5 scaling of four inflammation types as indicated. Bars in e-g represent mean values ±SD. One-way ANOVA followed by Tukey’s multiple comparison test was used. In b-f *=0.05 < p <0.01, **=0.01 < p < 0.001, ***=0.001 < p < 0.0001, ****=p < 0.0001. Panel **h** shows representative images of histopathological findings in lung sections of COH04S1-vaccinated animals and control animals. Black arrows indicate moderate and mild bronchioalveolar hyperplasia in lung sections of sMVA-IM and sMVA-IN control animals as well as COH04S1-IN animals. Black arrows in lung sections of unvaccinated control animals indicate hyperplastic alveolar cells. 10x magnification.

At day 10 post challenge, hamsters were euthanized for viral load assessment and histopathology analysis. Viral load was measured in the lungs and nasal turbinates/wash by quantification of SARS-CoV-2 genomic RNA (gRNA) and sub-genomic RNA (sgRNA) to gauge the magnitude of total and replicating virus at lower and upper respiratory tracts. Compared to lung viral loads of control animals, markedly reduced gRNA and sgRNA copies were observed in the lungs of COH04S1-IM and COH04S1-IN-vaccinated animals (Figs. 2c-d), demonstrating potent vaccine protection against lower respiratory tract infection through IM and IN vaccination. SARS-CoV-2 gRNA copies in the lungs of COH04S1-IM and COH04S1-IN-vaccinated animals were more than three to four orders of magnitude lower than in the lungs of control animals. Furthermore, while 10^3^-10^6^ sgRNA copies were detected in the lungs of control animals, sgRNA was undetectable in the lungs of COH04S1-IM and COH04S1-IN-vaccinated animals, indicating complete absence of replicating virus in lung tissue of vaccinated hamsters. Compared to nasal viral loads of controls, gRNA and sgRNA viral loads in nasal turbinates and wash of COH04S1-IM and COH04S1-IN-vaccinated animals appeared only marginally reduced, indicating limited vaccine impact on upper respiratory tract infection by COH04S1 independent of the vaccination route (Fig. S2).

Histopathological findings in hematoxylin/eosin-stained lung sections of euthanized animals were assessed by a board-certified pathologist and scored in a blinded manner on a scale from 1 to 5 based on the severity and diffusion of the lesions (Table S2). Control animals demonstrated compromised lung structure characterized by moderate bronchioloalveolar hyperplasia with consolidation of lung tissue, minimal to mild mononuclear or mixed cell inflammation, and syncytial formation (Figs. 2e-h). In contrast, COH04S1-IM-vaccinated animals did not show lung pathology of any type and grade in 6/6 hamsters, demonstrating potent vaccine protection against SARS-CoV-2-mediated lung injury in hamsters by IM vaccination with COH04S1. While COH04S1-IN-vaccinated animals presented no severe histopathological findings and significantly reduced lung pathology compared to controls, COH04S1-IN-vaccinated hamsters consistently showed a mild form of bronchioalveolar hyperplasia and grade 1 interstitial inflammation in a subset of animals, indicating that IN vaccination with COH04S1 mediated potent but incomplete vaccine protection against SARS-CoV-2-mediated lung damage in this model (Fig. 2e-h).

Correlative analysis of pre-challenge immunity and post-challenge outcome revealed that any of the evaluated COH04S1-induced responses, including S, RBD, and N-specific antibodies and NAb, correlated with protection from weight loss, lung infection, and lung pathology (Fig. S3), confirming that the observed protection was vaccine-mediated. These results in sum demonstrate that hamsters vaccinated IM and IN with COH04S1 are protected from progressive weight loss, lower respiratory tract infection, and severe lung pathology following SARS-CoV-2 challenge.

### 2D and 1D vaccination of NHP with COH04S1 stimulates robust antigen-specific binding antibodies, NAb responses, and antigen-specific IFNγ and IL-2 expressing T cells

NHP represent a mild COVID-19 disease model that is widely used to bolster preclinical SARS-CoV-2 vaccine efficacy against upper and lower respiratory tract infection in an animal species that is more closely related to humans^49–56^. We used the African green monkey NHP model to assess COH04S1 vaccine protection against SARS-CoV-2 by 2D and 1D vaccination regimens, referred to as COH04S1-2D and COH04S1-1D, respectively. For 2D vaccination, NHP were vaccinated twice in a four week interval with 2.5×10^8^ PFU of COH04S1. For 1D vaccination, NHP were vaccinated once with 5×10^8^ PFU of COH04S1 (Fig. 3a). As controls, NHP were either mock-vaccinated or vaccinated with empty sMVA vector via the same schedule and dose vaccination regimen. Robust serum binding antibodies to S, RBD, and N were stimulated in NHP by both COH04S1-2D and COH04S1-1D, whereas binding antibodies in the 2D vaccine group were strongly boosted after the second dose. At the time of challenge, S- and RBD-specific antibody titers measured in the 2D and 1D vaccine groups were comparable (Fig. 3b-d, and S4), while N-specific titers appeared higher in the 2D vaccine group than in the 1D vaccine group. Notably, similar S-specific antibody titers were measured in 2D- and 1D-vaccinated NHP by S antigens based on the original Wuhan-Hu-1 reference strain and various VOC (Beta, Gamma, Delta), whereas binding antibodies measured with the VOC-specific S antigens tended to be lower than those measured with the Wuhan-Hu-1 S antigen (Figs. 3e, and S4). Antigen-specific binding antibodies were also measured in lung bronchioalveolar lavage (BAL) samples two weeks pre-challenge. BAL IgG binding antibodies to S, RBD, and N were detected in both COH04S1-2D and COH04S1-1D-vaccinated NHP, although BAL IgG antibody titers measured at this time point pre-challenge were higher in the 2D vaccine group than in the 1D vaccine group (Figs. 3f, and S4).

**Figure 3.**
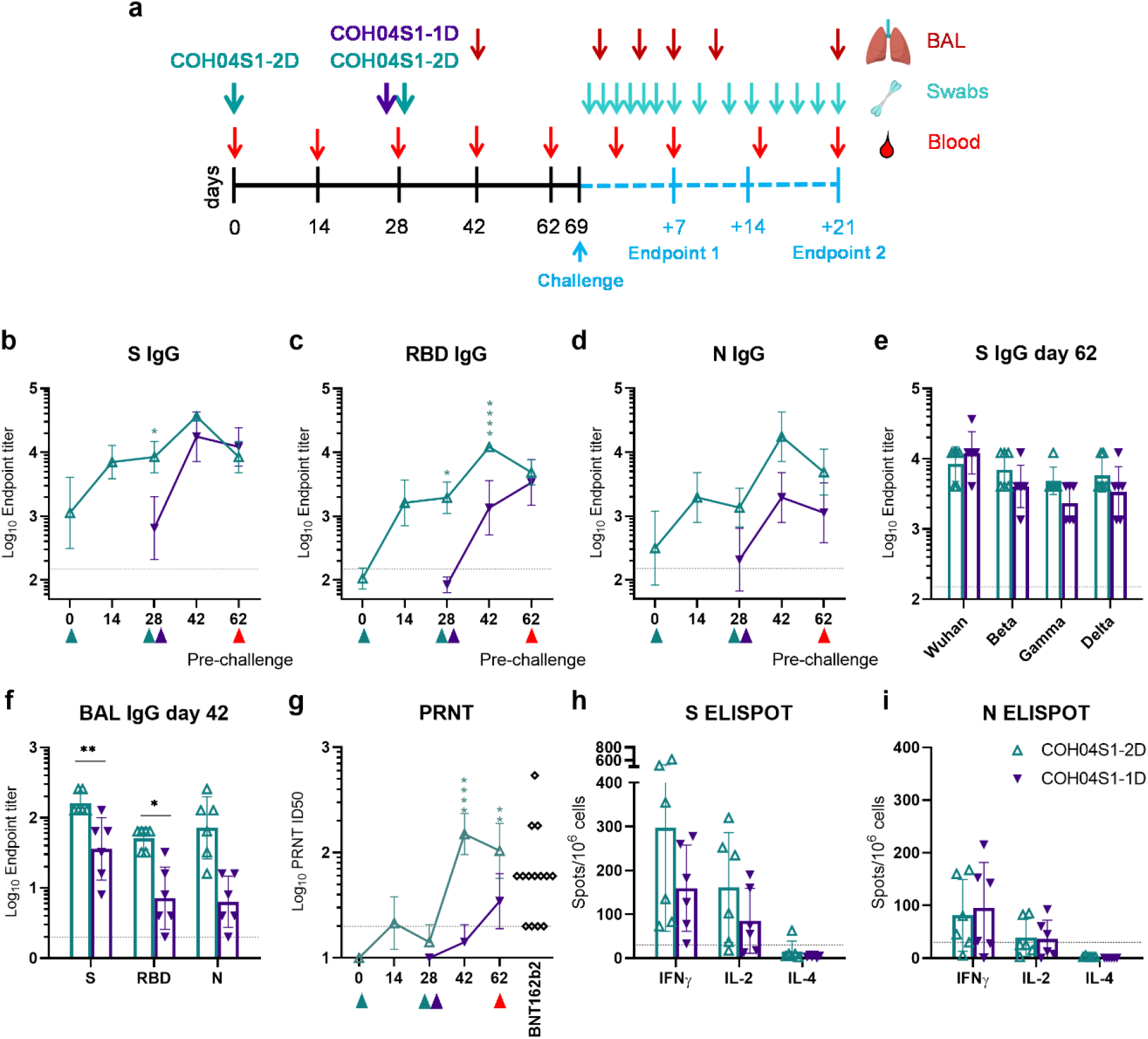
COH04S1 vaccine immunogenicity in NHP. **a. Study design.** African green monkey NHP were vaccinated with COH04S1 vaccine using two-dose (COH04S1-2D; N=6)) or single-dose (COH04S1-1D; N=6) immunization regimens as indicated. Mock-vaccinated NHP and NHP vaccinated with empty sMVA vector using the same dose immunization regimens were included as controls. Six weeks after immunization, NHP were IN/IT challenged with SARS-CoV-2 USA-WA1/2020 reference strain. Bronchoalveolar lavages (BAL) and oral and nasal swab samples for viral load analysis were collected at multiple time points post-challenge. A subset of animals in each group were necropsied at day 7 and 21 post-challenge for viral load and histopathology analysis. **b-d. Binding antibody titers**. S, RBD, and N antigen-specific binding antibody titers were measured at the indicated time points in serum samples of COH04S1-2D and COH04S1-1D-vaccinated NHP via ELISA. Data is presented as geometric mean values ± geometric SD. Two-way ANOVA followed by Sidak’s multiple comparison test was used. **e. VOC-specific antibody titers**. S-specific binding antibody titers were measured at day 62 by ELISA using S antigens based on the Wuhan-Hu-1 reference strain or several SARS-CoV-2 VOC, including Beta (B.1.351), Gamma (P.1), and Delta (B.1.617.2). Endpoint titers are presented as geometric mean values ± geometric SD. Two-way ANOVA with Sidak’s multiple comparison test was used for statistical analysis. **f. BAL antibody titers.** S, RBD, and N antigen-specific binding antibody titers were measured by ELISA at day 42 in BAL samples of COH04S1-IM and COH04S1-IN vaccine group. Endpoint titers are presented as geometric mean values ± geometric SD. Two-way ANOVA with Sidak’s multiple comparison test was used. **g. NAb titers**. NAb titers were measured by PRNT assay against SARS-CoV-2 infectious virus in serum samples of COH04S1-vaccinated NHP. Serum dilutions reducing the plaque number by 50% (ID50) are presented as geometric mean values ± geometric SD. Two-way ANOVA followed by Sidak’s multiple comparison test was used to compare ID50 values. PRNT-assayed NAb titers measured in serum samples of healthcare workers (N=14) vaccinated two times with Pfizer-BioNTech BNT162b2 mRNA vaccine were included. **h-i. T cell responses.** IFNγ, IL-2, and IL-4-expressing S and N-antigen-specific T cell responses were measured at day 42 by ELISPOT. Bars represent mean values, lines represent ±SD. Two-way ANOVA followed by Tukey’s multiple comparison test was used. Dotted lines indicate the arbitrary positive threshold of 30 spots/10^6^ cells. *=0.05 < p <0.01, **=0.01 < p < 0.001, ***=0.001 < p < 0.0001, ****=p < 0.0001.

NAb measurements based on PRNT assay revealed that both COH04S1-2D and COH04S1-1D elicited antibodies with efficacy to neutralize SARS-CoV-2 infectious virus (USA-WA1/2020). NAb responses measured in COH04S1-2D-vaccinated animals were boosted after the second dose and exceeded those measured in COH04S1-1D-vaccinated NHP at the time of challenge (Fig. 3g). Notably, NAb measured by PRNT assay in COH04S1-2D- and COH04S1-1D-vaccinated NHP were within the range of peak titers measured post-second dose in a cohort of healthcare workers that received the FDA-approved BNT162b2 mRNA vaccine from Pfizer/BioNTech.

At two weeks pre-challenge, we also assessed SARS-CoV-2-specific T cells in COH04S1-vaccinated NHP. Both 2D and 1D vaccination with COH04S1 stimulated robust IFNγ and IL-2-expressing S and N antigen-specific T cells, whereas no or only very low levels of IL-4-expressing S and N-specific T cells were observed in COH04S1-2D- or COH04S1-1D-vaccinated animals, consistent with Th1-biased immunity (Figs. 3h-i, and S5). S-specific T cells were generally detected at a higher frequency than N-specific T cells in both the COH04S1-2D and COH04S1- 1D vaccine groups. In addition, S-specific T cells measured at this time point pre-challenge were detected at a higher frequency in 2D-vaccinated NHP than in 1D-vaccinated NHP. These results demonstrate that COH04S1 elicits robust antigen-specific binding antibodies, NAb, and antigen-specific IFNγ and IL-2-expressing T cells in NHP through 2D and 1D vaccination regimens.

### 2D and 1D vaccination with COH04S1 protect NHP from lower and upper respiratory tract infection following SARS-CoV-2 challenge

Six weeks after 2D or 1D vaccination with COH04S1, NHP were challenged by IN/Intratracheal (IT) route with 1×10^5^ PFU of SARS-CoV-2 (USA-WA1/2020). SARS-CoV-2 viral loads in lower and upper respiratory tracts were assessed at several time points for 21 days post challenge in BAL and nasal/oral swabs by quantification of gRNA and sgRNA and infectious virus titers (Figs. 4a-i, 5a-i and S6). Progressively declining viral loads were overall measured over the 21 days post challenge observation period in BAL and nasal/oral swabs of both COH04S1-vaccinated NHP and control animals. Compared to BAL viral loads of control animals, markedly reduced gRNA and sgRNA copies and infectious virus titers were measured in BAL of COH04S1-2D and COH04S1-1D-vaccinated NHP at almost all time points post challenge (Figs. 4a-b, d-e, g-h), indicating a potent vaccine impact on lower respiratory tract infection. Notably, BAL gRNA and sgRNA copies and infectious titers measured in COH04S1-2D and COH04S1-1D-vaccinated NHP at day 2 immediately after challenge were on average 2-3 orders of magnitude lower than those measured in controls, confirming rapid vaccine efficacy. The marked reduction in BAL viral loads of COH04S1-vaccinated NHP compared to controls was confirmed when combining viral loads measured at all timepoints post challenge by area under the curve (AUC; Figs. 4c, f, i).

**Figure 4.**
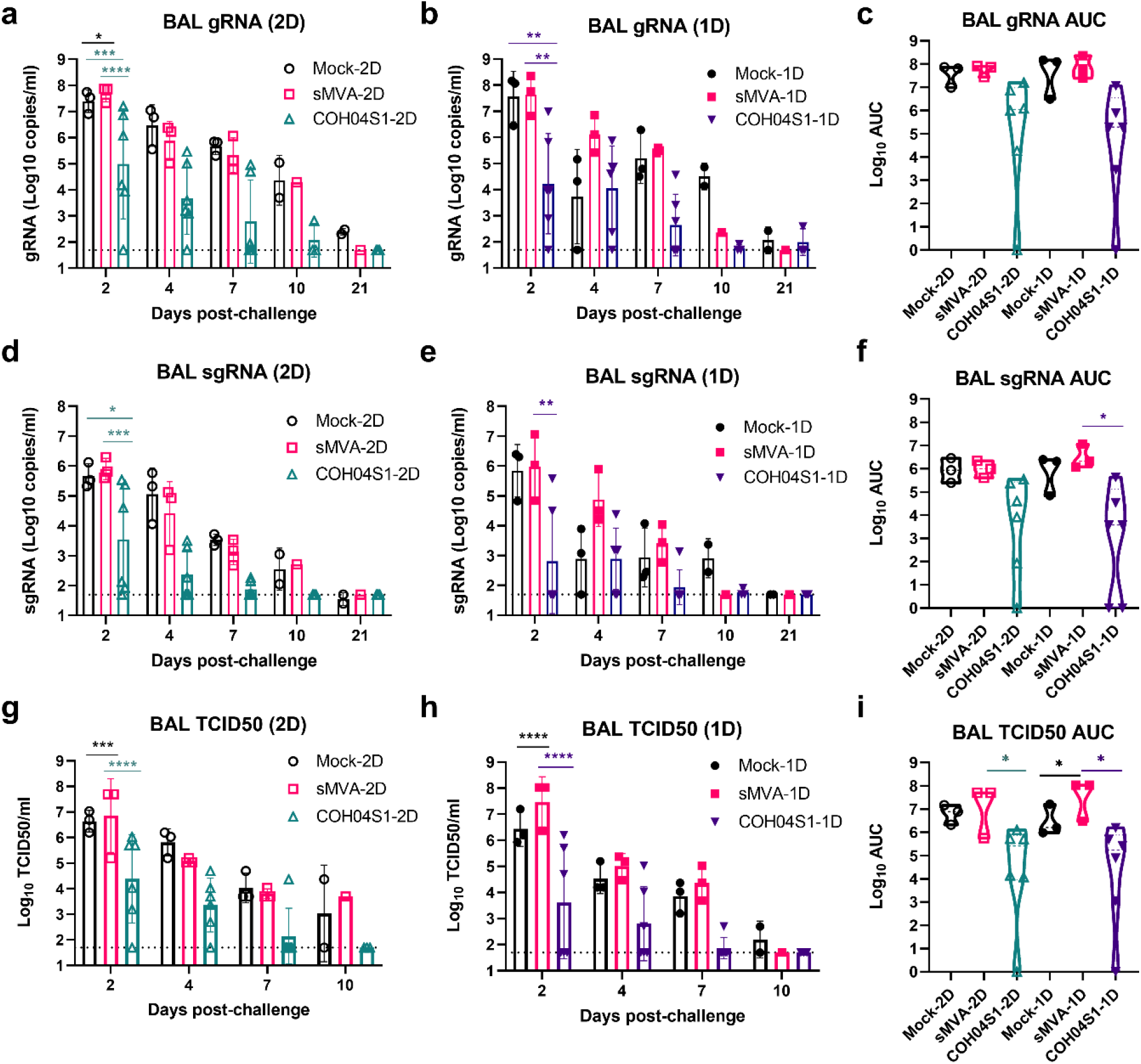
BAL viral loads in COH04S1-vaccinated NHP following SARS-CoV-2 challenge. SARS-CoV-2 genomic RNA (gRNA; a-c) and subgenomic RNA (sgRNA; d-f) copies and TCID50 infectious virus titers (g-i) were measured at the indicated days post challenge in bronchoalveolar lavages (BAL) of COH04S1-2D and COH04S1-1D-vaccinated NHP and mock-vaccinated and sMVA vector control animals. Bars represent geometric means, lines represent ±geometric SD. Dotted lines represent lower limit of detection. Two-way ANOVA followed by Sidak’s multiple comparison test was used. Panels **c**, **f**, and **i** show viral loads by area under the curve (AUC). Violin plots show values ranges with median values (dashed line) and quartiles (dotted line). AUC=0 was indicated as 1. One-way ANOVA followed by Tukey’s multiple comparison test was used. *=0.05 < p <0.01, **=0.01 < p < 0.001, ***=0.001 < p < 0.0001, ****=p < 0.0001.

**Figure 5.**
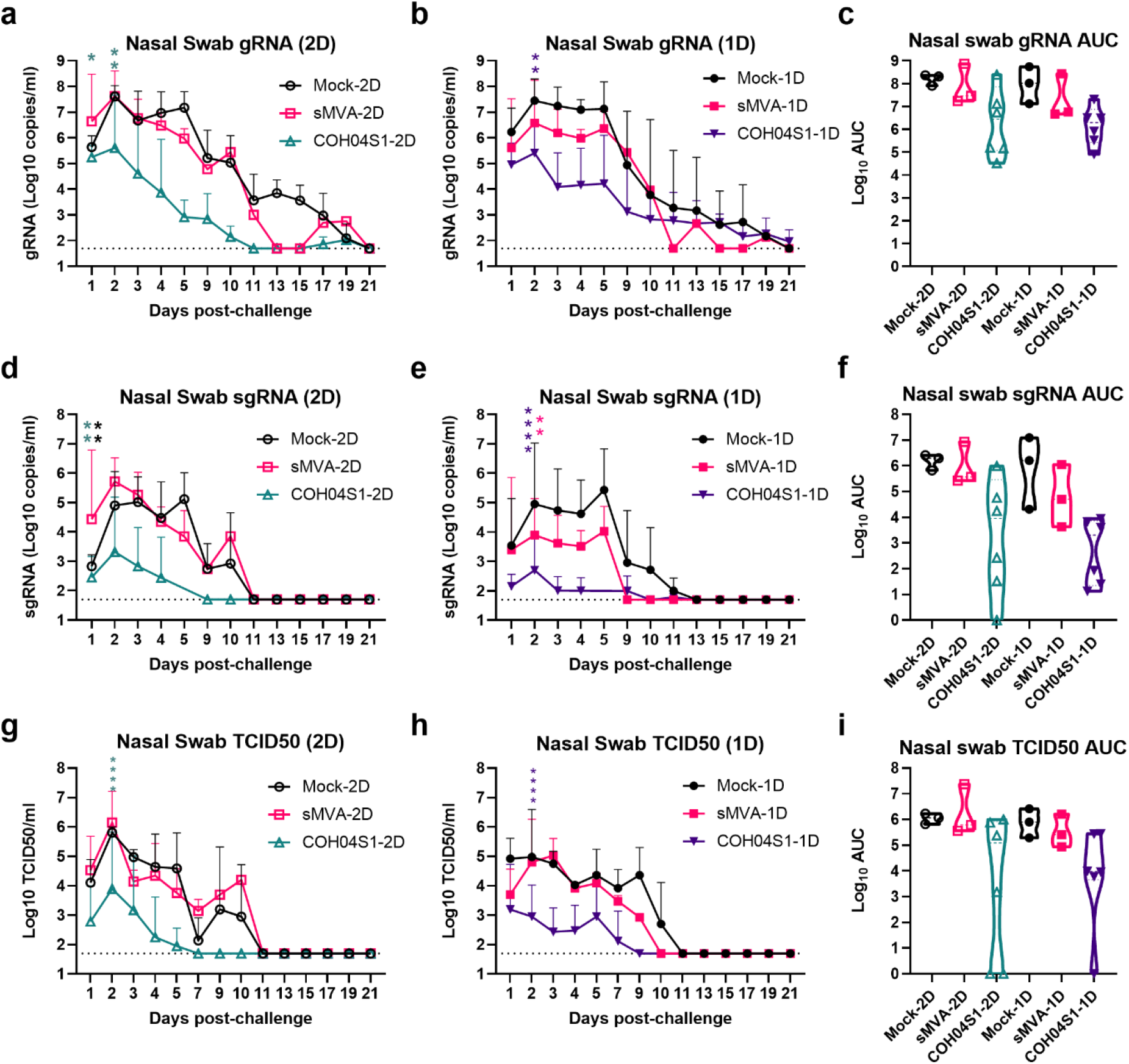
Nasal swab viral loads in COH04S1-vaccinated NHP after SARS-CoV-2 challenge. SARS-CoV-2 genomic RNA (gRNA; a-c) and subgenomic RNA (sgRNA; d-f) copies and TCID50 infectious virus titers (g-i) were measured at the indicated days post challenge in nasal swab samples of COH04S1-2D and COH04S1-1D-vaccinated NHP and mock-vaccinated and sMVA vector-vaccinated control animals. Lines represent geometric means + geometric SD. Dotted lines represent lower limit of detection. Two-way ANOVA followed by Sidak’s multiple comparison test was used. Panels **c**, **f**, and **i** show viral loads by area under the curve (AUC). Violin plots show values ranges with median values (dashed line) and quartiles (dotted line). AUC=0 was indicated as 1. One-way ANOVA followed by Tukey’s multiple comparison test was used. *=0.05 < p <0.01, **=0.01 < p < 0.001, ***=0.001 < p < 0.0001, ****=p < 0.0001.

Similar to viral loads in BAL of COH04S1-vaccinated NHP, gRNA and sgRNA copies and infectious virus titers measured at the first 10 days post challenge in nasal swabs of COH04S1-2D or COH04S1-1D-vaccinated NHP were consistently lower than those of control animals, demonstrating vaccine efficacy to prevent upper respiratory tract infection (Figs. 5a-i). In addition, gRNA and sgRNA in oral swabs of COH04S1-vaccinated NHP tended to be consistently lower than those in oral swabs of control animals (Fig. S6). Notably, nasal and oral swab gRNA and sgRNA copies and nasal swab infectious titers measured at 1-3 days immediately after challenge in COH04S1-vaccinated NHP were significantly reduced compared to those of controls, indicating immediate vaccine protection. Overall reduced nasal and oral swab viral loads in COH04S1-vaccinated NHP compared to control animals were confirmed when evaluating the nasal and oral swab viral loads over time by AUC (Figs. 5 c, f, i, and S6). No significant differences in viral loads were observed between COH04S1-2D and COH04S1-1D-vaccinated animals, indicating similar protection against lower and upper respiratory tract infection by COH04S1 through 2D and 1D vaccination. Viral loads measured at day 7 or 21 post challenge in lung tissue and several other organs of necropsied animals did not indicate evident differences between COH04S1-vaccinated NHP and control animals (Fig. S7). While gRNA and sgRNA copies measured at day 7 post challenge in lung samples of COH04S1-2D and COH04S1-1D-vaccinated NHP appeared reduced compared to those measured in lung samples of controls, these results were inconclusive due to low or undetectable gRNA and sgRNA levels in lung samples of the 1D mock-vaccinated control monkey (Fig. S7). Histopathological findings assessed by a board-certified pathologist in a blinded manner at day 7 and 21 post challenge were generally only minor and comparable between COH04S1 vaccine and control groups, and no increase in inflammation was observed for COH04S1-vaccinated NHP compared to control animals (Table S3). A strong inverse correlation could be assessed between vaccine-induced humoral and cellular immune responses, including S, RBD, and N-specific binding antibodies in serum and BAL, NAb, and S- and N-specific T cells, and SARS-CoV-2 viral loads in BAL and nasal swab samples (Fig. S8). These results in sum demonstrate that 2D or 1D vaccination regimens of COH04S1 protect NHP from lower and upper respiratory tract infection following IN/IT challenge with SARS-CoV-2.

### NHP vaccinated with 2D and 1D vaccination regimens of COH04S1 develop robust post-challenge anamnestic antiviral immune responses

To assess the vaccine impact on post-challenge viral immunity, humoral and cellular responses were evaluated in COH04S1-vaccinated NHP and control animals at 1, 2, and 3 weeks post-challenge (Figs. 6, S9). This analysis revealed that control animals developed robust binding antibodies to S, RBD, and N at 15-or 21-days post challenge (Figs. 6a-f), which indicated stimulation of potent humoral responses by the SARS-CoV-2 challenge virus in naïve NHP. In contrast, NAb titers measured post challenge in control animals by PRNT assay against infectious virus were in general relatively low or at the limit of detection (Figs. 6g, j), although elevated NAb responses were observed in 1D mock-vaccinated control animals at day 15 and 21 post challenge (Fig. 6j). In addition, no or only low level T cell responses were detected post-challenge in control animals, with the exception of elevated frequencies of S and N-specific IFNγ-expressing T cell responses in mock-vaccinated control animals at day 7 post challenge (Figs. 6h, k, i).

**Figure 6.**
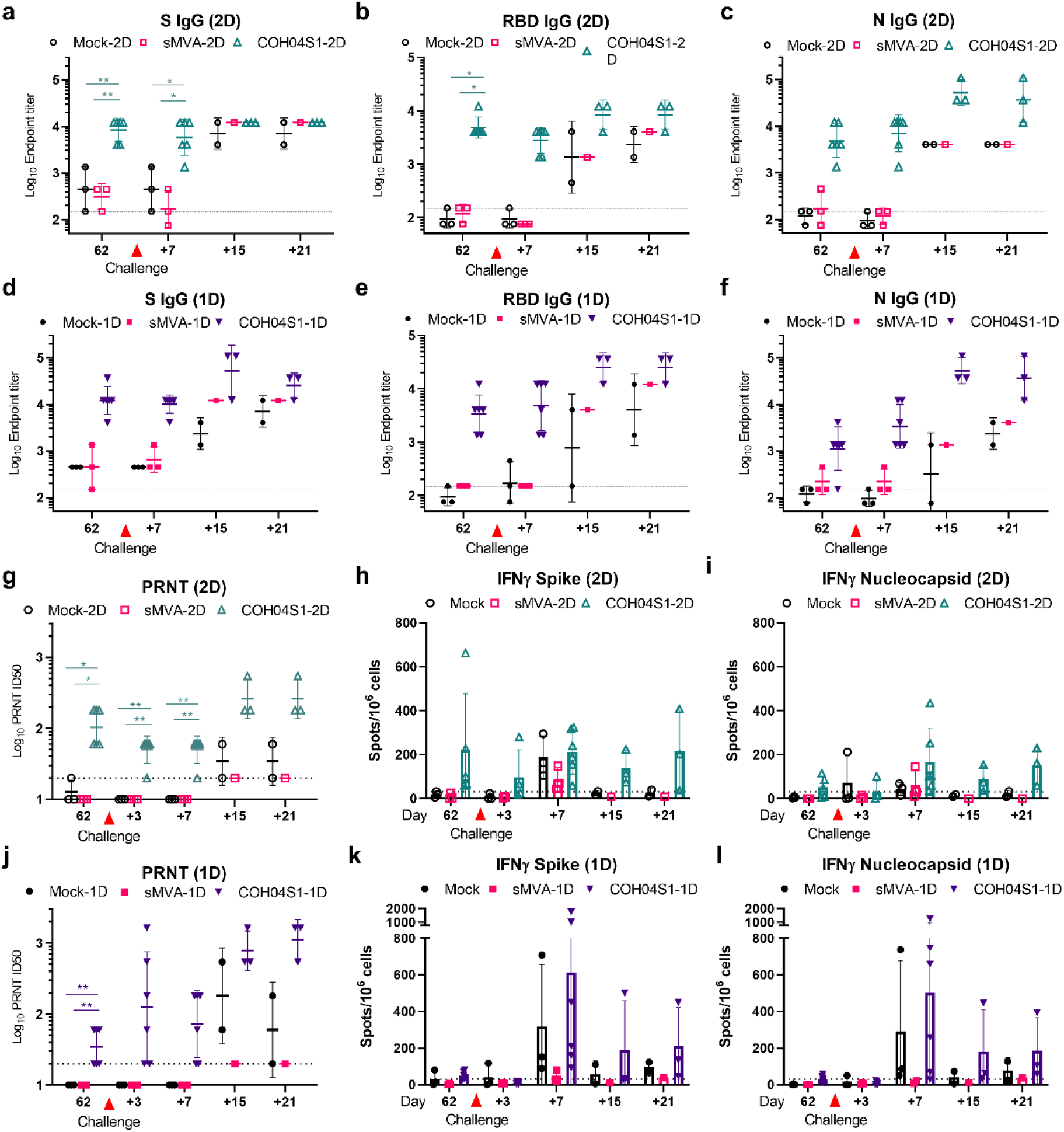
SARS-CoV-2-specific post-challenge immune responses in COH04S1-vaccinated NHP. SARS-CoV-2-specific humoral and cellular immune responses were measured at day 3, 7, 15 and 21 post-challenge in COH04S1-2D and COH04S1-1D-vaccinated NHP and mock-vaccinated and sMVA vector control animals and compared to pre-challenge immunity assessed in these groups. Panels **a-f** show S, RBD and N antigen specific binding antibody titers evaluated by ELISA. Panels **g-j** show NAb titers measured by PRNT assay against SARS-CoV-2 USA-WA1/2020 reference strain. Lines represent geometric means ± geometric SD. Dotted lines indicate lower limit of detection (LOD). Values below the LOD are indicated as ½ LOD. Panels **h-I, k-l** show IFNγ-expressing S and N-specific T cell responses measured by Fluorospot assay. Bars represent mean values, lines represent ±SD. Dotted lines indicate the arbitrary positive threshold of 30 spots/10^6^ cells. Two-way ANOVA followed by Tukey’s multiple comparison test was used. Time-points with <3 samples/group (d+15 and d+21) were excluded from the statistical evaluation. *0.05 < p <0.01, **0.01 < p < 0.001, ***0.001 < p < 0.0001, ****p < 0.0001.

Compared to antibodies measured pre-challenge in COH04S1-vaccinated NHP, boosted titers of S, RBD, and N antigen-specific binding antibodies and PRNT-assayed NAb were measured in COH04S1-vaccinated animals at day 15 and 21 post challenge (Figs. 6a-g, j), indicating induction of robust post-challenge anamnestic immune responses. This post-challenge booster effect on SARS-CoV-2-specific humoral immunity appeared more pronounced in COH04S1-1D-vaccinated NHP than in COH04S1-2D-vaccinated NHP, which may be a result of the lower pre-challenge responses in the 1D vaccine group compared to the 2D vaccine group. In addition, binding antibodies and NAb measured at day 15 and 21 post-challenge in COH04S1-vaccinated NHP generally exceeded those measured in control animals, indicating heightened vaccine-mediated humoral recall responses against SARS-CoV-2 through 2D or 1D vaccination. While S and N antigen-specific IFNγ-expressing T cell levels measured in COH04S1-2D-vaccinated animals tended to increase only slightly after challenge, a marked increase in post-challenge S- and N-specific IFNγ-expressing T cells was observed in COH04S1-1D-vaccinated NHP, indicating potent vaccine-mediated cellular recall responses following 1D vaccination. S and N-specific IL- 4-expressing T cells were either only very low or undetectable in COH04S1-vaccinated NHP and control animals at any time point post challenge, consistent with Th-1-biased immunity following challenge (Fig. S9). Correlation analysis did not unambiguously reveal a strong association between any of the post-challenge immunological parameters and post-challenge virological assessments (Fig. S8). These results demonstrate that NHP vaccinated with 2D or 1D vaccination regimens of COH04S1 develop potent post-challenge anamnestic antiviral responses.

## Discussion

The emergence of several SARS-CoV-2 VOC with the capacity to evade S-specific NAb threatens the efficacy of approved COVID-19 vaccines, which primarily utilize a single antigen design based solely on the S protein. One way to avoid or minimize the risk for SARS-CoV-2 evasion of vaccine-induced immunity could be the stimulation of broadly functional humoral and cellular immunity beyond the induction of S-specific NAb. Particularly the stimulation of T cells by a combination of multiple immunodominant antigens could act as an additional countermeasure to confer long-term and broadly effective immunity against SARS-CoV-2 and its emerging VOC^21–24^. Several recent studies indicate that SARS-CoV-2 VOC have the capacity to effectively escape humoral immunity, whereas they are unable to evade T cells elicited through natural infection and vaccination^57^.

Here, we demonstrate that multiantigenic sMVA-vectored SARS-CoV-2 vaccine COH04S1 co-expressing full-length S and N antigens provides potent immunogenicity and protective efficacy in animal models. We show in Syrian hamsters and NHP that COH04S1 elicits robust antigen-specific humoral and cellular immune responses and protects against SARS-CoV-2 challenge through different vaccination routes and dose regimens. While these animal studies were not designed to assess the contribution of N in COH04S1-mediated protective immunity, these results warrant further evaluation of COH04S1 in ongoing and future clinical trials. COH04S1 represents a second-generation COVID-19 vaccine candidate that could be used alone or in combination with other existing vaccines in parenteral or mucosal prime-boost or single-shot vaccination strategies to augment vaccine-mediated protective immune responses against SARS-CoV-2.

Although several SARS-CoV-2 vaccines based on MVA have been developed and evaluated in animal models for immunogenicity and efficacy^29,58–61^, there is currently no MVA-based SARS-CoV-2 vaccine approved for routine clinical use. COH04S1 and MVA-SARS-2-S, an MVA vector expressing S alone, are currently the only MVA-based SARS-CoV-2 vaccines that are clinically evaluated^61^. In addition, COH04S1 and a recently developed adenovirus vector approach are currently the only clinically evaluated SARS-CoV-2 vaccines that utilize an antigen combination composed of S and N^62^. These findings highlight the potential importance of COH04S1 as a second generation multiantigenic SARS-CoV-2 vaccine to contribute to the establishment of long-term protective immunity to prevent COVID-19 disease. In addition, these findings highlight the potential of the sMVA platform and synthetic biology in poxvirus-vectored vaccine technology to rapidly generate protective and clinical-grade vaccine vectors for infectious disease prevention.

COH04S1 demonstrated potent immunogenicity in Syrian hamsters by IM and IN vaccination and in NHP by 2D and 1D vaccination regimens to elicit robust SARS-CoV-2-specific humoral and cellular immune responses to both S and N antigens. This included high-titer S and N-specific binding antibodies in addition to robust binding antibodies targeting the RBD, the primary target of NAb. Binding antibodies elicited by COH04S1 in hamsters as well as T cell responses induced by COH04S1 in NHP indicated Th1-biased immunity, which is thought to be the preferred antiviral adaptive immune response to avoid vaccine-associated enhanced respiratory disease^63,64^. NAb elicited by COH04S1 in hamsters and NHP showed potent neutralizing activity against SARS-CoV-2 infectious virus, highlighting the potential of COH04S1 to induce antibody responses that are considered essential for protection against SARS-CoV-2. Notably, NAb titers stimulated by COH04S1 in NHP appeared similar to peak NAb titers stimulated in healthcare workers by two doses of the FDA approved Pfizer/BioNTech mRNA vaccine. In addition, NAb stimulated by COH04S1 in hamsters showed neutralizing activity against PV variants based on several SARS-CoV-2 VOC, including Alpha, Beta, Gamma, and the rapidly spreading Delta variant, indicating the capacity of COH04S1 to stimulate cross-protective NAb against SARS-CoV-2 VOC.

Both IM and IN vaccination with COH04S1 provided potent efficacy to protect Syrian hamsters from progressive weight loss, lower respiratory tract infection, and lung injury upon IN challenge with SARS-CoV-2, highlighting the potential of COH04S1 to stimulate protective immunity against respiratory disease through parenteral and mucosal vaccination routes. In contrast, IM or IN vaccination of hamsters with COH04S1 appeared to provide only limited protection against upper respiratory tract infection following viral challenge, indicating that COH04S1-mediated parenteral or mucosal immune stimulation afforded only little protection in this small animal model at the site of viral inoculation, which may have been associated with the relatively aggressive sub-lethal viral challenge dose. While IM and IN vaccination with COH04S1 provided similar immunogenicity and protective efficacy against SARS-CoV-2 in hamsters, IN vaccination with COH04S1 appeared superior compared to IM vaccination to protect against initial minor weight loss at the early phase after SARS-CoV-2 challenge. On the other hand, hamsters vaccinated IM with COH041S1 were completely protected from lung injury following challenge, while hamsters vaccinated IN with COH041S1 showed minor lung pathology and inflammation at day 10 post challenge, suggesting superior protection against viral- or immune-mediated lung pathology through IM vaccination compared to IN vaccination with COH04S1. Additional studies are needed to define the protection mediated by COH04S1 through IM and IN vaccination.

The immunogenicity and protective efficacy afforded by COH04S1 against SARS-CoV-2 in Syrian hamsters by IM and IN vaccination appears consistent with known properties of MVA-vectored vaccines. While MVA is well-known to stimulate robust immunity by IM vaccination, MVA has also been shown to elicit potent immunity through IN vaccination strategies at mucosal surfaces. A recombinant MVA vector expressing the S protein of Middle East Respiratory Syndrome coronavirus (MERS-CoV), a close relative of SARS-CoV-2, has recently been shown to be safe and immunogenic following IM administration in a Phase 1 clinical trial^65^. This MVA vaccine has also been shown to protect dromedary camels against MERS-CoV challenge by co-vaccination via IM and IN routes^66^. In addition, several animal studies have shown that IN vaccination with MVA vaccines is a potent stimulator of bronchus-associated lymphoid tissue (BALT), a tertiary lymphoid tissue structures within the lung that is frequently present in children and adolescents and that serves as a general priming site for T cells^67^. While the precise mechanism and levels of protection afforded by COH04S1 against SARS-CoV-2 in the Syrian hamster model by IM and IN vaccination remains unclear, especially at early phase after challenge, these findings support the use of COH04S1 to elicit SARS-CoV-2 protective immunity by mucosal vaccination.

In addition to the protection afforded by IM and IN vaccination with COH04S1 in hamsters, 2D and 1D vaccination with COH04S1 in NHP provided potent protection against lower and upper respiratory tract infection upon IN/IT challenge with SARS-CoV-2. These findings demonstrate protective efficacy of COH04S1 by different dose vaccination regimens in a larger animal model that is thought to be critical to assess preclinical vaccine efficacy. While 2D and 1D vaccination of NHP with COH04S1 stimulated similar S and RBD-specific binding antibodies at the time of viral challenge, 2D vaccination appeared to be more effective than 1D vaccination in stimulating humoral and cellular immune responses, including NAb. Despite these lower pre-challenge immune responses in 1D-vaccinated animals compared to 2D-vaccinated NHP, 1D and 2D vaccination of NHP with COH04S1 afforded similar protection against lower and upper respiratory tract infection following viral challenge, suggesting that a single shot with COH04S1 is sufficient to induce protective immunity to SARS-CoV-2. In addition, while 2D vaccination with COH04S1 appeared to elicit more potent pre-challenge responses than 1D vaccination, overall more potent post-challenge anamnestic antiviral immune responses were observed in 1D-vaccinated NHP compared to 2D-vaccinated NHP, suggesting that a single shot with COH04S1 is sufficient to effectively prime vaccine-mediated protective recall responses to SARS-CoV-2. Whether the protection afforded by COH04S1 in NHP through 2D and 1D vaccination regimens is primarily mediated by NAb or complemented by a combination of multiple S and N antigen-specific humoral and cellular effector functions remains to be determined in future studies^10^.

These results in sum demonstrate that COH04S1 has the capacity to elicit potent and cross-reactive Th1-biased S and N-specific humoral and cellular responses that protect hamsters and NHP from SARS-CoV-2 by different vaccination routes and dose regimens. This multi-antigen sMVA-vectored SARS-CoV-2 vaccine could be complementary to available vaccines to induce robust and long-lasting protective immunity against SARS-CoV-2 and its emerging VOC.

## Methods

### COH04S1 and sMVA vaccine vectors

COH04S1 is a double-plaque purified virus isolate derived from the previously described sMVA-N/S vector (NCBI Accession# MW036243), which was generated using the three-plasmid system of the sMVA platform and fowlpox virus TROVAC as a helper virus for virus reconstitution^29^. COH04S1 co-expresses full-length S and N antigen sequences based on the Wuhan-Hu-1 reference strain (NCBI Accession# NC_045512)^29^. Sequence identity of COH04S1 seed stock virus was assessed by PaqBio long-read sequencing. COH04S1 and sMVA vaccine stocks for animal studies were produced using chicken embryo fibroblasts (CEF) and prepared by 36% sucrose cushion ultracentrifugation and virus resuspension in 1 mM Tris-HCl (pH 9). Virus stocks were stored at −80°C and titrated on CEF by plaque immunostaining as described^29^. Viral stocks were validated for antigen insertion and expression by PCR, sequencing, and Immunoblot.

### Animals, study design and challenge

In life portion of hamsters and NHP studies were carried out at Bioqual Inc. (Rockville, MD). Thirty 6-8 weeks old Syrian hamsters were randomly assigned to the groups, with 3 females and 3 males in each group. Hamsters were IM or IN vaccinated four weeks apart with 1×10^8^ PFU of COH04S1 or sMVA vaccine stocks diluted in phosphate-buffered saline (PBS). Two weeks post-booster vaccination animals were challenged IN (50µl/nare) with 3×10^4^ PFU (or 1.99×10^4^ Median Tissue Culture Infectious Dose, [TCID50]) of SARS-CoV-2 USA-WA1/2020 (BEI Resources; P4 animal challenge stock, NR-53780 lot no. 70038893). The stock was produced by infecting Vero E6-hACE2 cells (BEI Resources NR-53726) at low MOI with deposited passage 3 virus and resulted in a titer of 1.99×10^6^ TCID50/ml. Sequence identity was confirmed by next generation sequencing. Body weight was recorded daily for 10 days. Hamsters were humanely euthanized for lung samples collection. A total of 24 African green monkeys (*Chlorocebus aethiops;* 20 females and 4 males) from St. Kitts weighting 3-6 kg were randomized by weight and sex to vaccine and control groups. For 2D vaccination, NHP were either two times mock-vaccinated with PBS (N=3) in four weeks interval or vaccinated twice for weeks apart with 2.5×10^8^ PFU of COH04S1 (N=6) or sMVA control vector (N=3) diluted in PBS. For 1D vaccination, NHP were either one times mock-vaccinated (N=3) with PBS or vaccinated once with 5×10^8^ PFU of COH04S1 (N=6) or sMVA control vector (N=3) diluted in PBS. At six weeks post vaccination, NHP were challenged with 1×10^5^ TCID50 of SARS-CoV-2 USA-WA1/2020 strain diluted in PBS via combined IT (1 ml)/IN (0.5 ml/nare) route. Necropsy was performed 7 days and 21 days following challenge and organs were collected for gross pathology and histopathology. All animal studies were conducted in compliance with local, state, and federal regulations and were approved by Bioqual and City of Hope Institutional Animal Care and Use Committees (IACUC).

### ELISA binding antibody detection

SARS-CoV-2-specific binding antibodies in hamster and NHP samples were detected by indirect ELISA using purified S, RBD, and N proteins (Sino Biological 40589-V08B1, 40592-V08H, 40588-V08B), or Beta, Gamma, and Delta VOC-specific S proteins (Acro Biosystems SPN-C52Hk, SPN-C52Hg, SPN-C52He). 96-well plates were coated with 100ul/well of S, RBD, or N protein at a concentration of 1ug/ml in PBS and incubated overnight at 4°C. For binding antibody detection in hamsters serum, plates were washed 5X with wash buffer (0.05% Tween-20/ PBS), then blocked with 250ul/well of blocking buffer (0.5% casein/ 154mM NaCl/ 10mM Tris-HCl [pH 7.6]) for 2 hours at room temperature. After washing, 3-fold diluted heat-inactivated serum in blocking buffer was added to the plates and incubated 2 hours at room temperature. After washing, anti-Hamster IgG HRP secondary antibodies measuring total IgG(H+L), IgG_1_, or IgG_2_/IgG_3_ (Southern Biotech 6061-05, 1940-05, 1935-05) were diluted 1:1000 in blocking buffer and added to the plates. After 1 hour incubation, plates were washed and developed with 1 Step TMB-Ultra (Thermo Fisher 34029). The reaction was stopped with 1M H_2_SO_4_ and plates were immediately read on FilterMax F3 (Molecular Devices). For binding antibody detection in NHP serum, a similar protocol was used. Wash buffer was 0.1% Tween-20 in PBS, and blocking buffer was 1% casein/PBS for RBD and N antigen ELISA and 4% Normal Goat Serum/1% casein/PBS for S ELISA. For IgG quantification in NHP BAL samples 1% BSA/PBS was used as blocking and sample buffers. Goat anti-Monkey IgG (H+L) secondary antibody (ThermoFisher PA1-84631) was diluted 1:10,000. Endpoint titers were calculated as the highest dilution to have an absorbance >0.100.

### PRNT assay

NAb were measured by PRNT assay using SARS-CoV-2 USA-WA1/2020 strain (Lot # 080420-900). The stock was generated using Vero-E6 cells infected with seed stock virus obtained from Kenneth Plante at UTMB (lot # TVP 23156). Vero E6 cells (ATCC, CRL-1586) were seeded in 24-well plates at 175,000 cells/well in DMEM/ 10% FBS/Gentamicin. Serial 3-fold serum dilutions were incubated in 96-well plates with 30 PFU of SARS-CoV-2 USA-WA1/2020 strain for 1 hour at 37°C. The serum/virus mixture was transferred to Vero-E6 cells and incubated for 1 hour at 37°C. After that, 1 ml of 0.5% methylcellulose media was added to each well and plates were incubated at 37°C for three days. Plates were washed, and cells were fixed with methanol. Crystal violet staining was performed, and plaques were recorded. IC50 titers were calculated as the serum dilution that gave a 50% reduction in viral plaques in comparison to control wells. Serum samples collected from City of Hope healthcare workers (N=14) at day 60 post Pfizer/BioNTech BNT162b2 mRNA vaccination were part of an IRB-approved observational study to establish durability of immunogenic properties of EUA vaccines against COVID-19 (IRB20720). Subjects gave informed consent.

### PV production

SARS-CoV-2 PV was produced using a plasmid lentiviral system based on pALD-gag-pol, pALD-rev, and pALD-GFP (Aldevron). Plasmid pALD-GFP was modified to express Firefly luciferase (pALD-Fluc). Plasmid pCMV3-S (Sino Biological VG40589-UT) was used and modified to express SARS-CoV-2 Wuhan-Hu-1 S with D614G modification. Customized gene sequences cloned into pTwist-CMV-BetaGlobin (Twist Biosciences) were used to express SARS-CoV-2 VOC-specific S variants (Table S1). All S antigen were expressed with C-terminal 19aa deletion. A transfection mixture was prepared 1ml OptiMEM that contained 30 µl of TransIT-Lenti transfection reagent (Mirus MIR6600) and 6 µg pALD-Fluc, 6 µg pALD-gag-pol, 2.4 µg pALD-rev, and 6.6 µg S expression plasmid. The mixture was added to 5×10^6^ HEK293T/17 cells (ATCC CRL11268) seeded the day before in 10 cm dishes and the cells were incubated for 72h at 37°C. Supernatant containing PV was harvested and frozen in aliquots at −80°C. Lentivirus was titrated using the Lenti-XTM p24 Rapid Titer Kit (Takara) according to the manufacturer’s instructions.

### PV neutralization assay

SARS-CoV-2 PV were titrated *in vitro* to calculate the virus stock amount that equals 100,000-200,000 relative luciferase units. Flat-bottom 96-well plates were coated with 100 μL poly-L-lysine (0.01%). Serial 2-fold serum dilutions starting from 1:20 were prepared in 50 μL media and added to the plates in triplicates, followed by 50 μL of PV. Plates were incubated overnight at 4°C. The following day, 10,000 HEK293T-ACE2 cells^68^ were added to each well in the presence of 3 µg/ml polybrene and plates were incubated at 37°C. After 48h of incubation, luciferase lysis buffer (Promega E1531) was added, and luminescence was quantified using SpectraMax L (Molecular Devices. 100 μL Luciferase Assay Reagent (Promega E1483) luciferin/well, 10 seconds integration time). For each plate, positive (PV only) and negative (cells only) controls were added. The neutralization titer for each dilution was calculated as follows: NT = [1−(mean luminescence with immune sera/mean luminescence without immune sera)] × 100. The titers that gave 50% neutralization (NT50) were calculated by determining the linear slope of the graph plotting NT versus serum dilution by using the next higher and lower NT using Office Excel (v2019).

### ELISPOT T cell detection

Peripheral blood mononuclear cells (PBMC) were isolated from fresh blood using Ficoll and counted using Luna-FL cell counter (Logos Biosystems). Pre-immune samples were evaluated using Human IFN-γ/IL-4 Double-Color FluoroSpot (ImmunoSpot); however, this kit only allowed to assess NHP IFN-γ but not NHP IL-4. The remaining time-points were evaluated using Monkey IFNγ/IL-4 FluoroSpot FLEX kit and Monkey IL-2 FluoroSpot FLEX kit (Mabtech, X-21A16B and X-22B) following manufacturer instructions. Briefly, 200,000 cells/well in CTL-test serum free media (Immunospot CTLT-010) were added to duplicate wells and stimulated with peptide pools (15-mers, 11 aa overlap, >70% purity). Spike peptide library (GenScript) consisted of 316 peptides and was divided into 4 sub-pools spanning the S1 and S2 domains (1S1=1-86; 1S2=87-168; 2S1=169-242; 2S2=243-316; peptides 173 and 304-309 were not successfully synthesized and were therefore excluded from the pools). Nucleocapsid (GenScript) and Membrane (in house synthesized) libraries consisted of 102 and 53 peptides, respectively. Each peptide pool (2 µg/ml) and αCD28 (0.1 µg/ml, Mabtech) were added to the cells and plates were incubated for 48h at 37°C. Control cells (25,000/well) were stimulated with PHA (10 µg/ml). After incubation, plates were washed, and primary and secondary antibodies were added according to manufacturer’s protocol. Fluorescent spots were acquired using CTL S6 Fluorocore (Immunospot). For each sample, spots in unstimulated DMSO-only control wells were subtracted from spots in stimulated wells. Total spike response was calculated as the sum of the response to each spike sub-pool.

### gRNA quantification

SARS-CoV-2 gRNA copies per ml nasal wash, BAL fluid or swab, or per gram of tissue were quantified by qRT-PCR assay (Bioqual, SOP BV-034) using primer/probe sequences binding to a conserved region of SARS-CoV-2 N gene. Viral RNA was isolated from BAL fluid or swabs using the Qiagen MinElute virus spin kit (57704). For tissues, viral RNA was extracted with RNA-STAT 60 (Tel-test B)/ chloroform, precipitated and resuspended in RNAse-free water. To generate a control for the amplification, RNA was isolated from SARS-CoV-2 virus stocks. RNA copies were determined from an O.D. reading at 260, using the estimate that 1.0 OD at A260 equals 40 µg/ml of RNA. A final dilution of 10^8^ copies per 3 µl was then divided into single use aliquots of 10 µl. These were stored at −80°C until needed. For the master mix preparation, 2.5 ml of 2X buffer containing Taq-polymerase, obtained from the TaqMan RT-PCR kit (Bioline BIO-78005), was added to a 15 ml tube. From the kit, 50 µl of the RT and 100 µl of RNAse inhibitor were also added. The primer pair at 2 µM concentration was then added in a volume of 1.5 ml. Lastly, 0.5 ml of water and 350 µl of the probe at a concentration of 2 µM were added and the tube vortexed. For the reactions, 45 µl of the master mix and 5 µl of the sample RNA were added to the wells of a 96-well plate. All samples were tested in triplicate. The plates were sealed with a plastic sheet. The control RNA was prepared to contain 10^6^ to 10^7^ copies per 3 µl. Eight 10-fold serial dilutions of control RNA were prepared using RNAse-free water by adding 5 µl of the control to 45 µl of water. Duplicate samples of each dilution were prepared as described. For amplification, the plate was placed in an Applied Biosystems 7500 Sequence detector and amplified using the following program: 48°C for 30 minutes, 95°C for 10 minutes followed by 40 cycles of 95°C for 15 seconds, and 1 minute at 55°C. The number of copies of RNA per ml was calculated by extrapolation from the standard curve and multiplying by the reciprocal of 0.2 ml extraction volume. This gave a practical range of 50 to 5 x 10^8^ RNA copies per swabs or per ml BAL fluid. Primer/probe sequences: 5’-GAC CCC AAA ATC AGC GAA AT-3’; 5’-TCT GGT TAC TGC CAG TTG AAT CTG-3’; and 5’-FAM-ACC CCG CAT TAC GTT TGG TGG ACC-BHQ1-3’

### sgRNA quantification

SARS-CoV-2 sgRNA copies were assessed through quantification of N gene mRNA by qRT-PCR using primer/probe sequences that were specifically designed to amplify and bind to a region of the N gene mRNA that is not packaged into virions. SARS-CoV-2 RNA was extracted from samples as described above. The signal was compared to a known standard curve of plasmid containing a cDNA copy of the N gene mRNA target region to give copies per ml. For amplification, the plate was placed in an Applied Biosystems 7500 Sequence detector and amplified using the following program: 48°C for 30 minutes, 95°C for 10 minutes followed by 40 cycles of 95°C for 15 seconds, and 1 minute at 55°C. The number of copies of RNA per ml was calculated by extrapolation from the standard curve and multiplying by the reciprocal of 0.2 ml extraction volume. This gave a practical range of 50 to 5 x 10^7^ RNA copies per swab or ml BAL fluid. Primer/probe sequences: 5’-CGA TCT CTT GTA GAT CTG TTC TC-3’; 5’-GGT GAA CCA AGA CGC AGT AT-3’; 5’-FAM-TAA CCA GAA TGG AGA ACG CAG TGG G-BHQ-3’

### Infectious titer quantification

Vero TMPRSS2 cells (Vaccine Research Center-NIAID) were plated at 25,000 cells/well in DMEM/10% FBS/Gentamicin. Ten-fold dilutions of the sample starting from 20 ul of material were added to the cells in quadruplicated and incubated at 37°C for 4 days. The cell monolayers were visually inspected, and presence of CPE noted. TCID50 values were calculated using the Read-Muench formula.

### Histopathology

Histopathological evaluation of hamsters and NHP lung sections were performed by Experimental Pathology Laboratories, Inc. (Sterling, VA) and Charles River (Wilmington, MA) respectively. At necropsy organs were collected and placed in 10% neutral buffered formalin for histopathologic analysis. Tissues were processed through to paraffin blocks, sectioned once at approximately 5 microns thickness, and stained with hematoxylin/eosin. Board certified pathologists were blinded to the vaccine groups and mock controls were used as a comparator.

### Statistical analyses

Statistical analyses were performed using Prism 8 (GraphPad, v8.3.0). One-way ANOVA with Holm-Sidak’s multiple comparison test, two-way ANOVA with Tukey’s or Dunn’s multiple comparison test, Kruskal-Wallis test followed by Dunn’s multiple comparison test, and Spearman correlation analysis were used. All tests were two-sided. The test applied for each analysis and the significance level is indicated in each figure legend. Prism 8 was used to derive correlation matrices.

## Acknowledgements

We would like to thank Angela Wong and Christina Ulloa for helping with laboratory logistics and secretarial support. We would like to acknowledge the Bioqual team for their technical support. We gratefully acknowledge City of Hope’s unique internal funding mechanism of selected scientific discoveries, enabling preclinical and clinical development to speedily advance products toward commercialization, such as COH04S1. The following reagent was deposited by the Centers for Disease Control and Prevention and obtained through BEI Resources, NIAID, NIH: SARS-Related Coronavirus 2, Isolate USA-WA1/2020, NR-53780. Don J. Diamond was partially supported by the following Public Health Service grants: U19 AI128913, CA111412, CA181045, and CA107399. The content is solely the responsibility of the authors and does not necessarily represent the official views of the National Institutes of Health.

## Author contributions

F.W. and F.C. supervised the study, designed the experiments, analyzed the data, and prepared the manuscript. D.J.D. designed the experiments, provided supervision, and oversaw final manuscript preparation. F.C., V.H.N., Y.P., H.C., V.K., K.F., J.N., M.K., D.J., J.M., A.I., Q.Z., and T.K. performed the *in vitro* experiments. S.K., A.S., H.A., and M.G.L. supervised clinical care of the animals and virological assays. P.F. assisted in statistical analyses. Y.S. provided managerial support. All the authors contributed to and approved the final version of this manuscript.

