## Supplementary information for "Synthetic Multiantigen MVA Vaccine COH04S1 Protects Against SARS-CoV-2 in Syrian Hamsters and Non-Human Primates"

**Table S1.** SARS-CoV-2 VOC-specific S mutations included in PV particles

| VOC | CDC Classification | Substitutions/Deletions (Δ) |
|---|---|---|
| B.1.1.7 | Alpha | Δ69/70, Δ144, N501Y, A570D, D614G, P681H, T716I, S982A, D1118H |
| B.1351 | Beta | L18F, D80A, D215G, Δ242-244, R246I, K417N, E484K, N501Y, D614G, A701V |
| P.1 | Gamma | L18F, T20N, P26S, D138Y, R190S, K417T, E484K, N501Y, D614G, H655Y, T1027I, V1176F |
| B.1.617.2 | Delta | T19R, G142D, Δ156-157, R158G, A222V, L452R, T478K, D614G, P681R, D950N |

**Table S2.** Scoring parameters used to evaluate hamster lung histopathology

| Grade | Severity | Findings |
|---|---|---|
| 1 | Minimal | histopathologic change ranging from inconspicuous to barely noticeable but so minor, small, or infrequent as to warrant no more than the least assignable grade. For multifocal or diffusely-distributed lesions, this grade was used for processes where less than approximately 10% of the tissue in an average high-power field was involved. For focal or diffuse hyperplastic/hypoplastic/ atrophic lesions, this grade was used when the affected structure or tissue had undergone a less than approximately 10% increase or decrease in volume. |
| 2 | Mild | histopathologic change that is a noticeable but not a prominent feature of the tissue. For multifocal or diffusely-distributed lesions, this grade was used for processes where between approximately 10% and 25% of the tissue in an average high-power field was involved. For focal or diffuse hyperplastic/hypoplastic/atrophic lesions, this grade was used when the affected structure or tissue had undergone between an approximately 10% to 25% increase or decrease in volume. |
| 3 | Moderate | histopathologic change that is a prominent but not a dominant feature of the tissue. For multifocal or diffusely-distributed lesions, this grade was used for processes where between approximately 25% and 50% of the tissue in an average high-power field was involved. For focal or diffuse hyperplastic/hypoplastic/atrophic lesions, this grade was used when the affected structure or tissue had undergone between an approximately 25% to 50% increase or decrease in volume. |
| 4 | Marked | histopathologic change that is a dominant but not an overwhelming feature of the tissue. For multifocal or diffusely-distributed lesions, this grade was used for processes where between approximately 50% and 95% of the tissue in an average high-power field was involved. For focal or diffuse hyperplastic/hypoplastic/atrophic lesions, this grade was used when the affected structure or tissue had undergone between an approximately 50% to 95% increase or decrease in volume. |
| 5 | Severe | histopathologic change that is an overwhelming feature of the tissue. For multifocal or diffusely-distributed lesions, this grade was used for processes where greater than approximately 95% of the tissue in an average high-power field was involved. For focal or diffuse hyperplastic/hypoplastic/atrophic lesions, this grade was used when the affected structure or tissue had undergone a greater than approximately 95% increase of decrease in volume. |

**Table S3.** NHP vaccine groups, necropsy schedule, and gross pathological findings

| Group | Study | Weight (kg) | Sex | Nx day +7 | Nx day +21 | Gross pathology findings at Nx |
|---|---|---|---|---|---|---|
| Mock | 2D | 4.76 | F | | X | All lobes normal |
| | | 3.02 | F | X | | Right caudal lung, remaining lobes normal. Overall hemorrhaging |
| | | 3.78 | F | | X | Left and Right caudal lung, remaining lobes normal |
| | 1D | 3.56 | F | X | | Pathology and congestion in all lobes |
| | | 3.45 | F | | X | Pathology and congestion in all lobes |
| | | 6.79 | M | | X | Left and Right caudal lung, remaining lobes normal |
| sMVA | 2D | 4.03 | F | X | | Right caudal lung, remaining lobes normal |
| | | 3.31 | F | X | | Gross congestion, possibly due to BALs. Lobes normal |
| | | 3.63 | F | | X | Right caudal lung, remaining lobes normal |
| | 1D | 3.41 | F | X | | Gross congestion, possibly due to BALs. Lobes normal |
| | | 3.73 | F | X | | Gross congestion, possibly due to BALs. Lobes normal |
| | | 6.05 | M | | X | All lobes normal |
| COH04S1 | 2D | 4.02 | F | X | | Gross congestion, possibly due to BALs. Lobes normal |
| | | 2.91 | F | X | | Right caudal lung, remaining lobes normal |
| | | 3.69 | F | | X | Gross congestion, possibly due to BALs. Lobes normal |
| | | 3.68 | F | | X | Right caudal lung, remaining lobes normal |
| | | 3.37 | F | X | | Right caudal lung, remaining lobes normal |
| | | 7.23 | M | | X | Right caudal lung, remaining lobes normal |
| | 1D | 3.79 | F | X | | Gross congestion, possibly due to BALs. Lobes normal |
| | | 3.83 | F | | X | Right caudal lung, remaining lobes normal |
| | | 2.95 | F | X | | Gross congestion, possibly due to BALs. Lobes normal |
| | | 3.86 | F | | X | Gross congestion, possibly due to BALs. Lobes normal |
| | | 3.36 | F | X | | Right caudal lung, remaining lobes normal |
| | | 5.63 | M | | X | All lobes normal |

### Figure S1

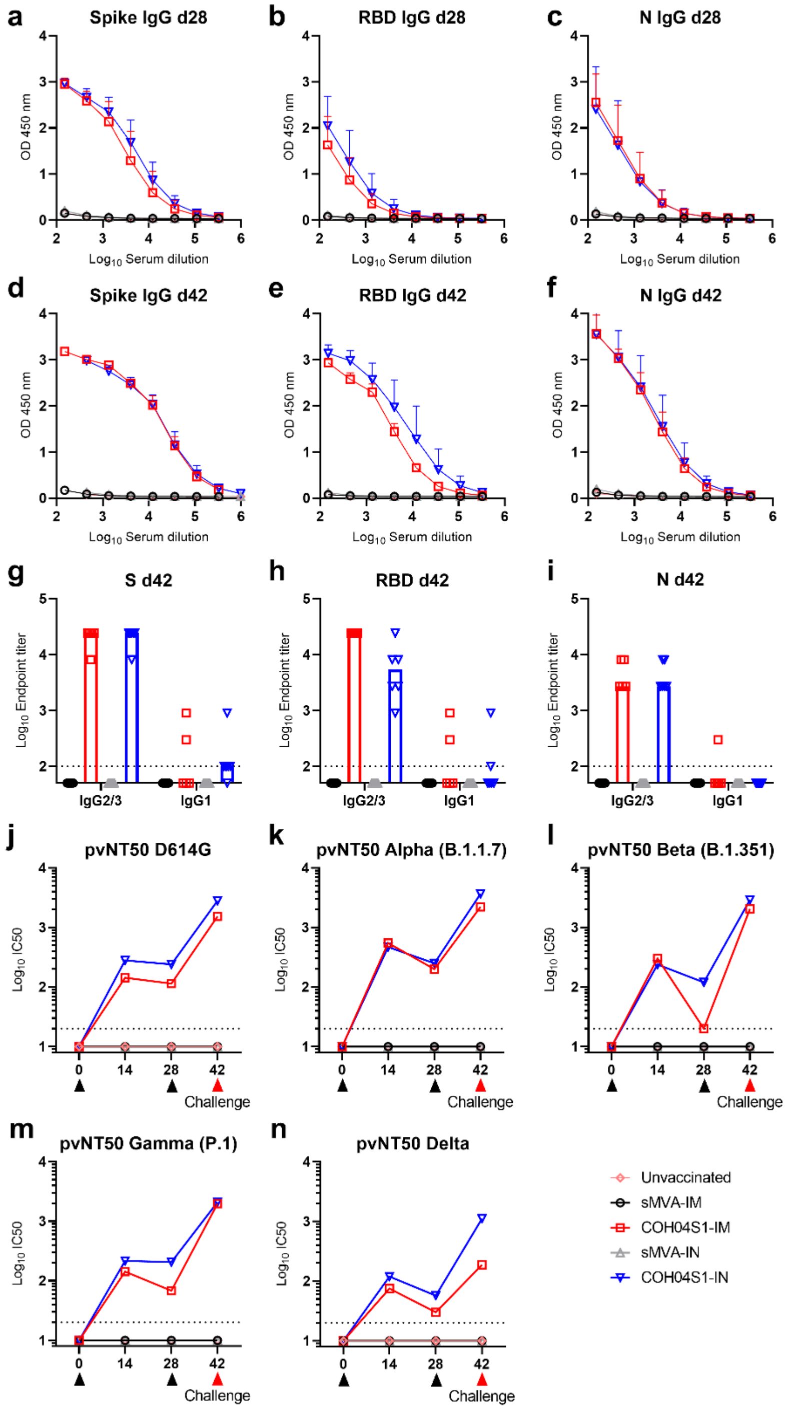


**Figure S1. Related to Figure 1. COH04S1 immunogenicity in Syrian hamsters. a-f. Binding antibody titers.** Shown are binding curves of S, RBD, and N antigen-specific antibody titers measured at day 28 (d28) or day 42 (d42) after the first and second vaccination in serum samples of COH04S1-IM and COH04S1-IN-vaccinated hamsters and unvaccinated and sMVA vector control animals. Two-way ANOVA with Tukey's multiple comparison test was used **g-I. IgG2-**

**3/IgG1 antibody titers.** Shown are IgG endpoint titers of S, RBD, and N antigen-specific IgG2-3 and IgG1 binding antibodies measured in serum samples of day 42 (d42). Geometric means are indicated with a line. Dotted lines indicate lower limit of detection. **j-n. VOC-specific NAb titers.** Shown are NAb titers measured in vaccine and control groups at the indicated time points by PV variants with D614G mutation or mutations based on several SARS-CoV-2 VOC.

**Figure S2**

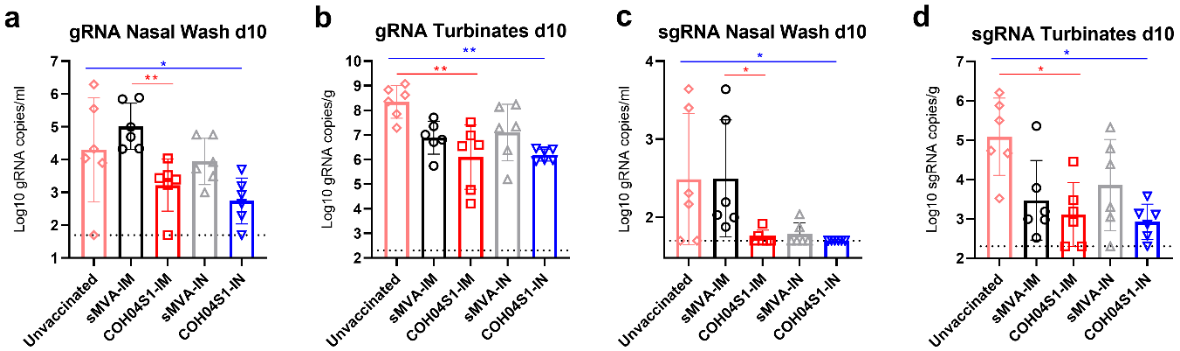


**Figure S2. Related to Figure 2. COH04S1-mediated vaccine protection of hamsters following sub-lethal SARS-CoV-2 challenge.** Shown are SARS-CoV-2 genomic RNA (gRNA) and sub-genomic RNA (sgRNA) copies measured by qPCR in nasal wash and nasal turbinates of COH04S1-IM and COH04S1-IN-vaccinated hamsters and unvaccinated and sMVA control animals at day 10 post-challenge. Bars show RNA copies geometric mean ± geometric SD. Dotted lines represent lower limit of detection of the assay. Kruskal-Wallis test followed by Dunn's multiple comparison test was used. *=0.05 < p <0.01, **=0.01 < p < 0.001

### Figure S3

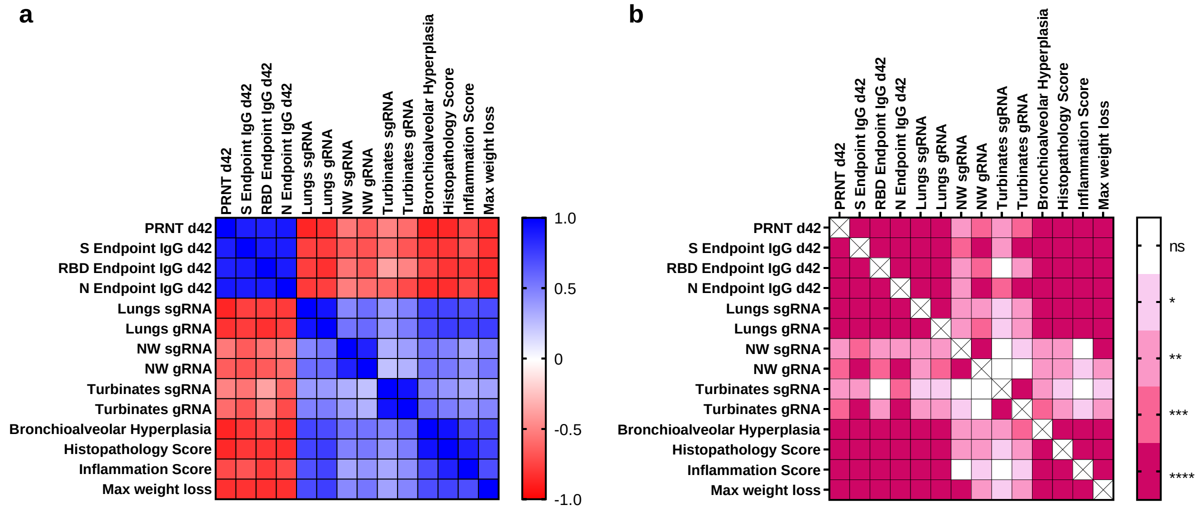


**Figure S3. Related to Figures 1-2. Correlative analysis of immunological and virological parameters in COH04S1-vaccinated hamsters challenged with SARS-CoV-2.** A Spearman correlation analyses was performed between the indicated pre-challenge immune responses and post-challenge virological assessments of COH04S1-IM and COH04S1-IN-vaccinated hamsters and unvaccinated and sMVA vector control animals. **a.** Spearman correlation coefficients were calculated and plotted as a matrix. **b.** Two-tailed p values were calculated and indicated as: *=0.05 < p <0.01, **=0.01 < p < 0.001, ***=0.001 < p < 0.0001, ****=p < 0.0001. ns= not significant.

### Figure S4

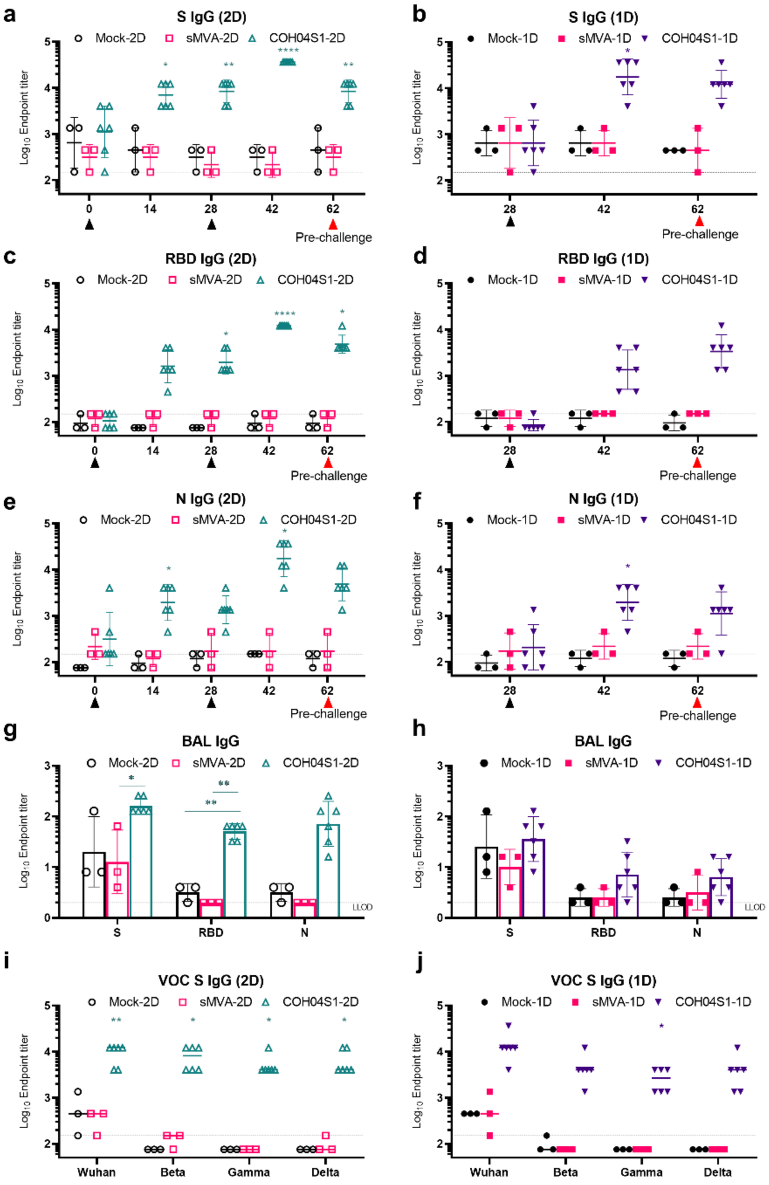


**Figure S4. Related to Figure 3. COH04S1-induced humoral immunity in NHP. a-f. Binding antibody titers.** Shown are S, RBD, and N antigen-specific IgG endpoint titers measured at the indicated time points of the study (Figure 3a) in COH04S1-2D and COH04S1-1D-vaccinated NHP and mock-vaccinated and sMVA vector control animals. Data are presented as geometric mean values ± geometric SD. Two-way ANOVA followed by Sidak's multiple comparison test was used. **g-h. BAL binding antibodies**. Shown are endpoint titers of IgG binding antibodies to S, RBD, and N measured in bronchoalveolar lavage (BAL) samples in vaccine and control groups at day 42 of the study (Figure 3a) two weeks after 2D or 1D vaccination. Endpoint titers are presented as geometric mean values ± geometric SD. Two-way ANOVA with Sidak's multiple comparison test was used. **i, j. VOC-specific antibody titers**. Shown are endpoint titers of S-specific IgG binding antibodies measured at day 62 of the study (Figure 3a) by ELISA with S antigens based on Wuhan-Hu-1 reference strain or several VOC, including Beta (B.1.351), Gamma (P.1), and Delta (B.1.617.2). Endpoint titers are presented as geometric mean values ± geometric SD. Two-way ANOVA with Sidak's multiple comparison test was used. *=0.05 < p <0.01, **=0.01 < p < 0.001, ***=0.001 < p < 0.0001, ****=p < 0.0001.

### Figure S5

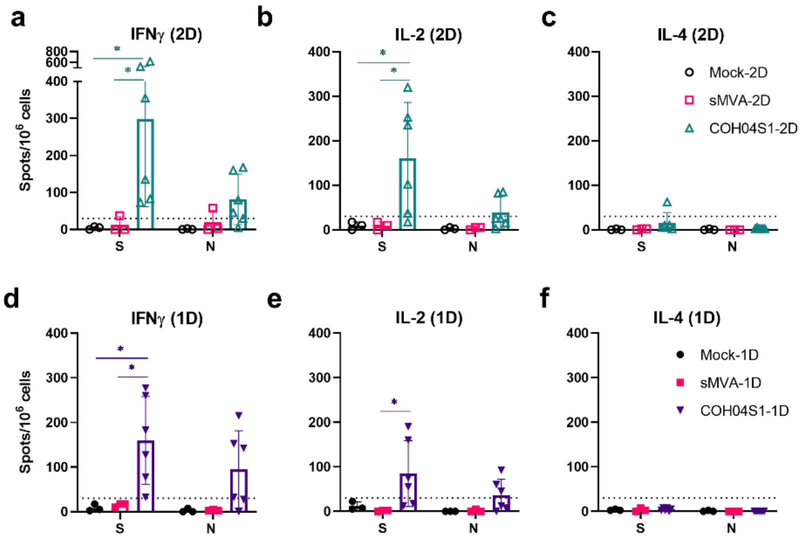


**Figure S5. Related to Figure 3. COH04S1-induced T cell responses in NHP. a-f.** Shown are IFNγ, IL-2, and IL-4-expressing S and N-specific T cell responses measured at day 42 of the study (Figure 3a) by Fluorospot assay in COH04S1-2D and COH04S1-1D-vaccinated NHP and mock-vaccinated and sMVA vector control animals. Bars represent mean values, lines represent ±SD. Two-way ANOVA followed by Tukey's multiple comparison test was used. Dotted lines indicate the arbitrary positive threshold of 30 spots/$10^6$ cells. *=0.05 < p <0.01.

### Figure S6

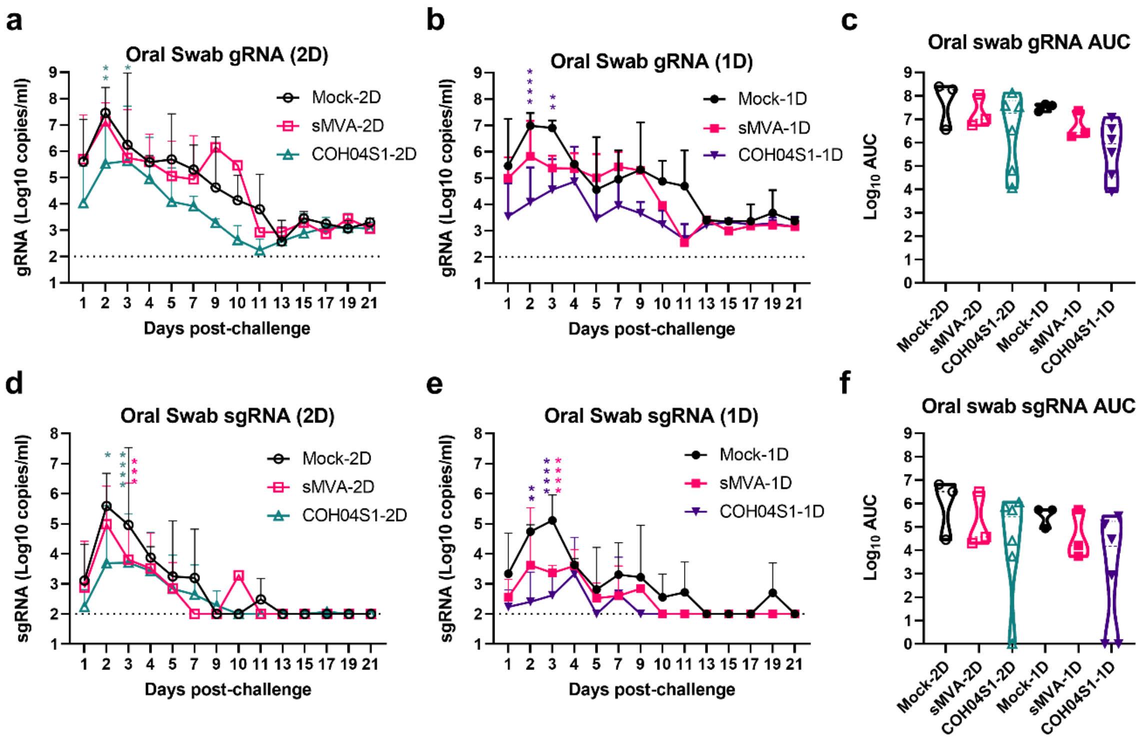


**Figure S6. Related to figure 4-5. Oral swab viral loads in COH04S1-vaccinated NHP following SARS-CoV-2 challenge.** SARS-CoV-2 genomic RNA (gRNA; a-c) and subgenomic RNA (sgRNA; d-f) copies were measured by qPCR at the indicated days post challenge in oral swab samples of COH04S1-2D and COH04S1-1D-vaccinated NHP and mock-vaccinated and sMVA vector control animals. Lines geometric means ± geometric SD. Dotted lines represent assay lower limit of detection. Two-way ANOVA followed by Sidak's multiple comparison test was used. Panels in **c** and **f** show viral loads by area under the curve (AUC). Violin plots show values ranges with median values (dashed line) and quartiles (dotted line). AUC=0 was indicated as 1. One-way ANOVA followed by Tukey's multiple comparison test was used. *=0.05 < p <0.01, **=0.01 < p < 0.001, ***=0.001 < p < 0.0001, ****=p < 0.0001.

### Figure S7

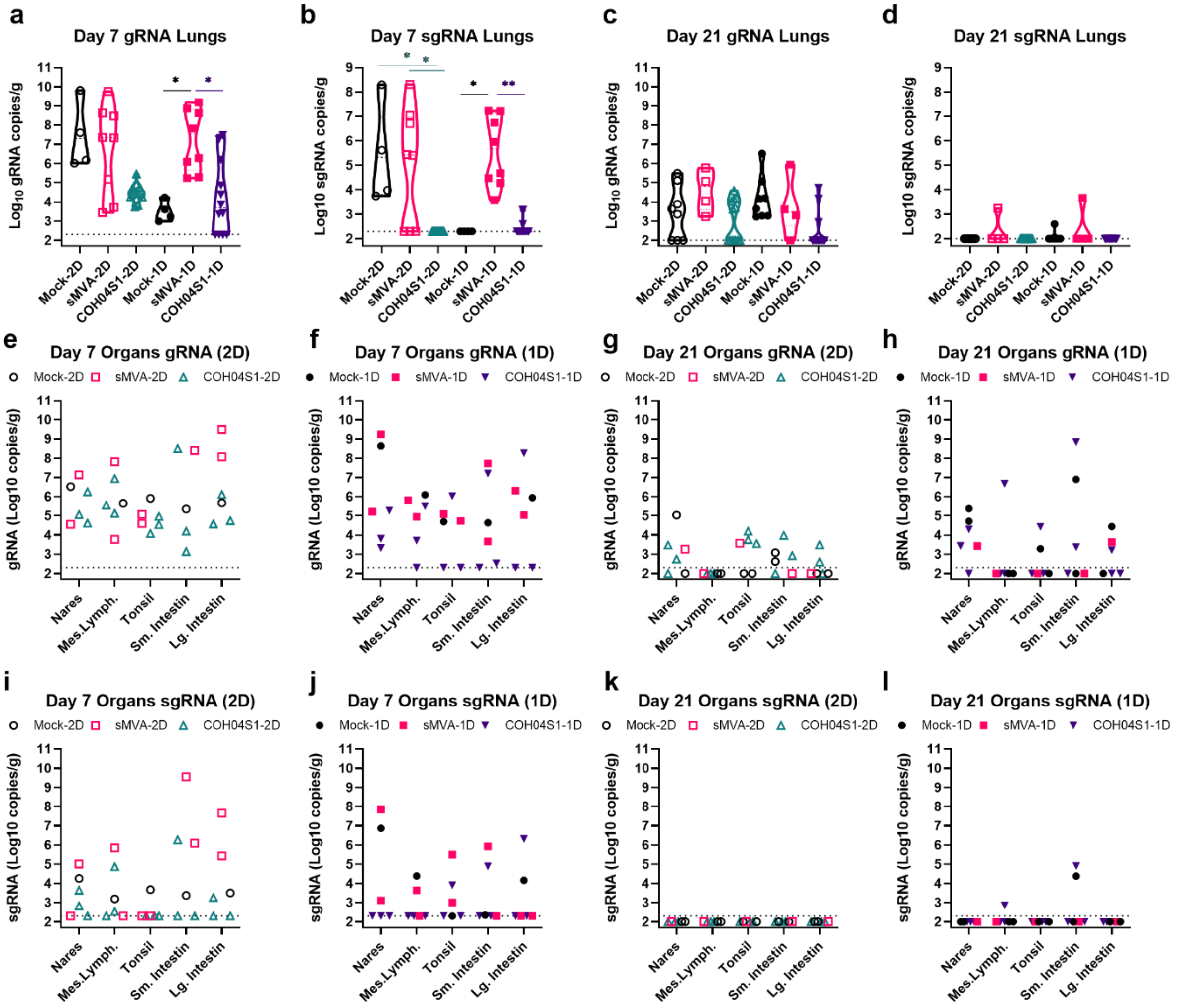


**Figure S7. Related to Figure 4-5. SARS-CoV-2 viral loads in lungs and other organs of COH04S1 vaccinated NHP following challenge.** Given are SARS-CoV-2 genomic RNA (gRNA; a-c) and subgenomic RNA (sgRNA; d-f) copies measured by qPCR at day 7 and 21 post challenge in the lungs and several other organs of necropsied COH04S1-2D and COH04S1-1D-vaccinated NHP and mock-vaccinated and sMVA vector control animals. Four lung samples were analyzed for each necropsied NHP (see Table S3 for necropsy schedule). Dotted lines represent assay lower limit of detection. Kruskal-Wallis test followed by Dunn's multiple comparison test was used. In e-l statistical evaluation was not performed because of the limited number of samples in the control groups. *=0.05 < p <0.01, **=0.01 < p < 0.001

**Figure S8**

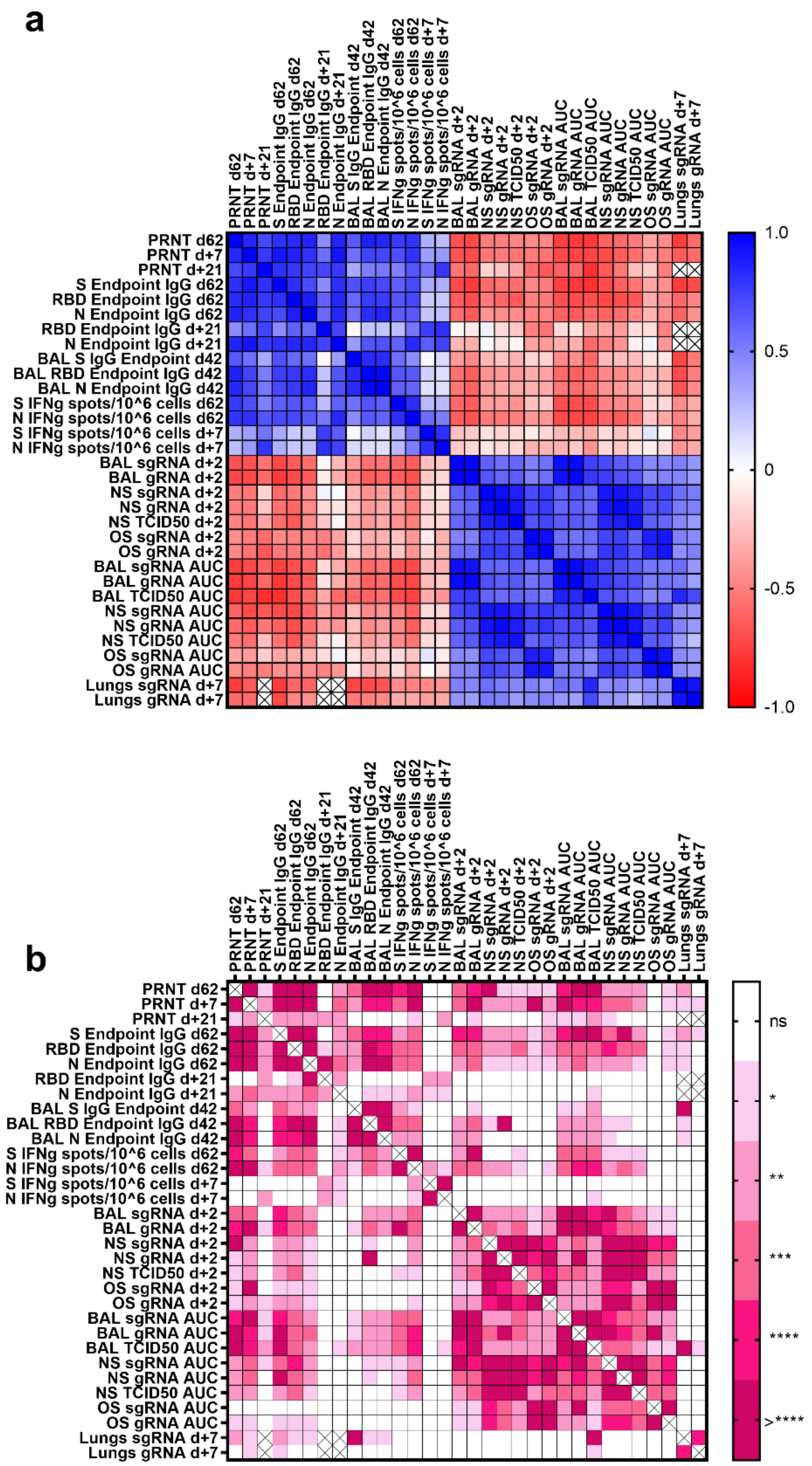


**Figure S8. Related to Figures 3-6. Correlative analysis of immunological and virological parameters in COH04S1-vaccinated NHP challenged with SARS-CoV-2.** Spearman correlation analysis was performed between the indicated pre-challenge and post-challenge immune responses and post-challenge virological assessments of COH04S1-2D and COH04S1-1D-vaccinated NHP and mock-vaccinated and sMVA control animals. Post-challenge time-points are indicated with “+“ before the day number. **a.** Spearman correlation coefficients were calculated and plotted as a matrix. **b.** Two-tailed p values were calculated and indicated as: *=0.05 < p <0.01, **=0.01 < p < 0.001, ***=0.001 < p < 0.0001, ****=p < 0.0001. ns= not significant.

**Figure S9**

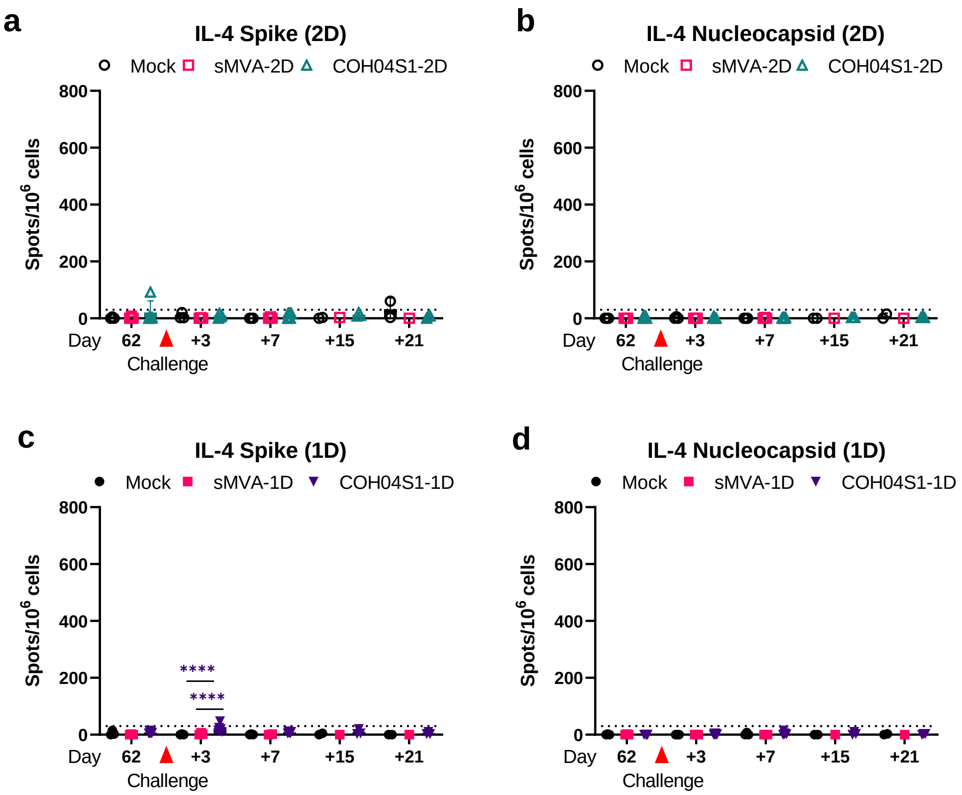


**Figure S9.** Related to **Figure 6. SARS-CoV-2-specific post-challenge cellular IL-4-specific responses in COH04S1-vaccinated NHP.** SARS-CoV-2-specific cellular immune responses were measured at day 3, 7, 15 and 21 post-challenge in COH04S1-2D and COH04S1-1D-vaccinated NHP and mock-vaccinated and sMVA vector control animals and compared to pre-challenge immunity assessed in these groups. IL-4-expressing S and N-specific T cell responses were measured by Fluorospot assay. Bars represent mean values, lines represent ±SD. Dotted lines indicate the arbitrary positive threshold of 30 spots/$10^6$ cells. Two-way ANOVA followed by Tukey's multiple comparison test was used. Time-points with <3 samples/group (d+15 and d+21) were excluded from the statistical evaluation. ****$p < 0.0001$.
